# Quantifying the mechanisms of vaccine-induced serotype replacement in *Streptococcus pneumoniae* using genome-informed modelling

**DOI:** 10.64898/2026.06.29.735370

**Authors:** Zein Assad, Kevin La, Corinne Levy, Robert Cohen, Assiya El Mniai, Inès Fafi, Zaba Valtuille, Stéphane Bechet, François Corrard, Andreas Werner, Christophe Rodriguez, Melissa N’Debi, Stephanie W. Lo, Lindsay Osei, Manon Jaboyedoff, François Angoulvant, Stéphane Bonacorsi, Philippe Bidet, André Birgy, Loïc de Pontual, Romain Basmaci, Emmanuelle Varon, Thibaut Morel-Journel, Naïm Ouldali

## Abstract

*Streptococcus pneumoniae* is a colonizer of the child’s nasopharynx and a leading cause of invasive pneumococcal disease (IPD). Despite widespread use of pneumococcal conjugate vaccines, recent rebounds in IPD incidence have coincided with serotype replacement. Whether this phenomenon is driven by the replacement of serotypes within lineages or the replacement of the lineages themselves is unclear. Here, we quantified the relative contribution of these two mechanisms to serotype replacement in carriage and their consequences for IPD. We combined nationwide longitudinal surveillance of carriage and IPD in children from 2002 to 2023 with whole-genome sequencing. Observed lineage-specific carriage rates across PCV-related periods were fitted with polynomial logistic regression and compared with two counterfactual scenarios, each isolating one mechanism with: (i) fixed serotype composition within varying lineages and (ii) serotype shifts within fixed lineage carriage rates over time. Carriage dynamics were consistent with an 82% contribution of serotype shifts within persistent lineages. Estimates of serotype-specific IPD incidence derived from this predominant scenario correlated with observed IPD trends. These findings quantify the dominant role of serotype replacement within persistent lineages, disentangling the vaccine-driven adaptive history of *S. pneumoniae*.

## Introduction

Vaccination reshapes host–pathogen interactions at both individual and population levels and imposes a selective force on microbial populations, conceptually analogous to antimicrobial pressure^1^. Although some vaccine-preventable pathogens can be driven toward elimination through interruption of transmission^2^, opportunistic pathogens – commensal bacteria capable of causing invasive disease – generally persist under vaccine pressure. In species such as *Haemophilus influenzae*, *Neisseria meningitidis*, and *Streptococcus pneumoniae*, antigenic variability and asymptomatic carriage within the respiratory microbiota maintain a persistent reservoir of diverse variants. By targeting only a subset of this diversity, vaccination drives population restructuring rather than eradication^3–5^.

*S. pneumoniae* colonizes the nasopharynx of young children, where asymptomatic carriage is a prerequisite for invasive pneumococcal disease (IPD) and constitutes the reservoir sustaining horizontal spread within the community^6^. The transition from carriage to disease is largely determined by the polysaccharide capsule, a defining surface structure that governs disease potential and immunogenicity. More than 100 antigenically distinct serotypes have been described, each differing in their propensity to cause disease, thereby linking carriage to IPD. Capsular polysaccharides have therefore been used as antigens in pneumococcal vaccines, which target the serotypes most frequently associated with IPD^7–10^.

Despite the widespread use of pneumococcal conjugate vaccines (PCVs), IPD remains a major cause of global morbidity and mortality, with over 300,000 deaths annually among children under five^11^. By inducing robust mucosal immunity, PCVs markedly reduce nasopharyngeal carriage of vaccine serotypes in vaccinated infants, and this decline generates strong herd protection by limiting transmission at the community level. Accordingly, the introduction of the seven-valent pneumococcal conjugate vaccine (PCV7), followed by PCV10 and PCV13, led to profound reductions in vaccine-type carriage and substantial decreases in IPD incidence^12^. However, PCVs do not target all serotypes, and the near elimination of vaccine serotypes from the nasopharynx has reshaped the pneumococcal population makeup: non-vaccine serotypes emerged to fill the ecological niche left vacant by the vaccine serotypes, while overall carriage prevalence in young children remained largely unchanged^13,14^. Serotype replacement has been reported across multiple countries and has contributed to a resurgence of IPD^15–17^. This led to continuous expansion of vaccine valency, including the recent implementation of next-generation formulations such as PCV15 and PCV20, with others in development^18^.

The substantial residual disease burden, despite continual escalation of vaccine valency, underscores the limits of serotype-based vaccination strategies and raises questions about the long-term sustainability of their impact. Indeed, currently implemented PCVs primarily target the serotypes of an obligate human commensal, thereby reshaping antigenic composition and enabling replacement by non-vaccine potentially invasive clones^19^. Within stable human reservoir, capsular serotype diversity – often attributed to diversifying selection by host immune responses – represents only one dimension of pneumococcal adaptation and vaccine escape. Genetic lineage reflects deeper ecological adaptation and offers insight into the mechanisms by which pneumococcal populations respond to vaccine-induced selective pressure^3^.

Over the past 15 years, analyses of pneumococcal population structure have shown that individual serotypes are frequently distributed across multiple genetically distinct lineages, underscoring the evolutionary flexibility of the species^20^. The recent development of the Global Pneumococcal Sequencing Project provides a unified, genome-wide lineage (defined as Global Pneumococcal Sequence Cluster, GPSC) classification, enabling robust comparisons of pneumococcal populations across geographic regions^21–23^. Based on genomic studies, two mechanisms of serotype replacement have been suggested: the expansion of non-vaccine-type lineages to occupy niches vacated by vaccine-type lineages^24^, and shifts in serotype composition within persistent lineages through the loss of vaccine serotypes and gain of non-vaccine serotypes^23,25^. However, the relative contribution of these mechanisms to the observed evolutionary dynamics is unknown.

Most genomic studies to date have focused on isolates from IPD^26–29^, lacking a complete view of pneumococcal diversity and evolution. As an opportunistic commensal, *S. pneumoniae* evolves not only under capsule-mediated immune selection but also through broader ecological interactions within the nasopharynx microbiota, including horizontal gene exchange^30^. A comprehensive understanding of pneumococcal dynamics therefore requires situating the organism within its ecological context rather than considering the disease landscape alone^13^. Although carriage studies have described shifts in serotype and lineage distributions across vaccine eras, and lineage-associated antimicrobial resistance and virulence genes, they have often been limited in scale or duration^31–34^. We postulate that disentangling the drivers of pneumococcal ecological dynamics requires integrating changes observed in carriage with their downstream consequences for invasive disease, through longitudinal analyses spanning multiple vaccine eras and jointly examining both compartments simultaneously and within the same population.

Here, we combined nationwide longitudinal surveillance of pneumococcal carriage and IPD within the same paediatric population to dissect the mechanisms underlying serotype replacement following the implementation of PCV7 and PCV13. Using genomic data, we constructed a reference model describing observed GPSC carriage dynamics while accounting for temporal structure and penicillin resistance. We then compared these estimates with two counterfactual scenarios – fixed serotype composition within lineages and fixed lineage carriage rates over time – to quantify the relative contributions of lineage turnover and within-lineage serotype replacement. Finally, we estimated serotype IPD incidence from scenario- derived carriage data to assess how these mechanisms reflect observed changes in invasive disease.

## Results

### Study population

#### Serotype data

Between 2002 and 2023 in France, 12,144 children aged ≤2 years with acute otitis media underwent nasopharyngeal sampling, from which *S. pneumoniae* was isolated in 7575 cases (overall carriage prevalence, 62.4%). Over the same period, 11,181 cases of IPD were estimated in children aged ≤17 years^35^, of which 6776 isolates were submitted to the National Reference Centre for Pneumococci (NRCP) for serotyping and antimicrobial susceptibility testing. These isolates span multiple vaccine eras, including the nationwide implementation of PCV7 from 2003 and PCV13 in 2010 (**Fig. 1** and Supplementary Results 1). Among all isolates, 74 distinct serotypes were identified, comprising 15% PCV7 serotypes, 26.2% additional PCV13 serotypes not included in PCV7 (PCV13 non-PCV7 serotypes), and 58.8% non-PCV13 serotypes (of which 2.0% were non-capsulated).

**Fig 1.**
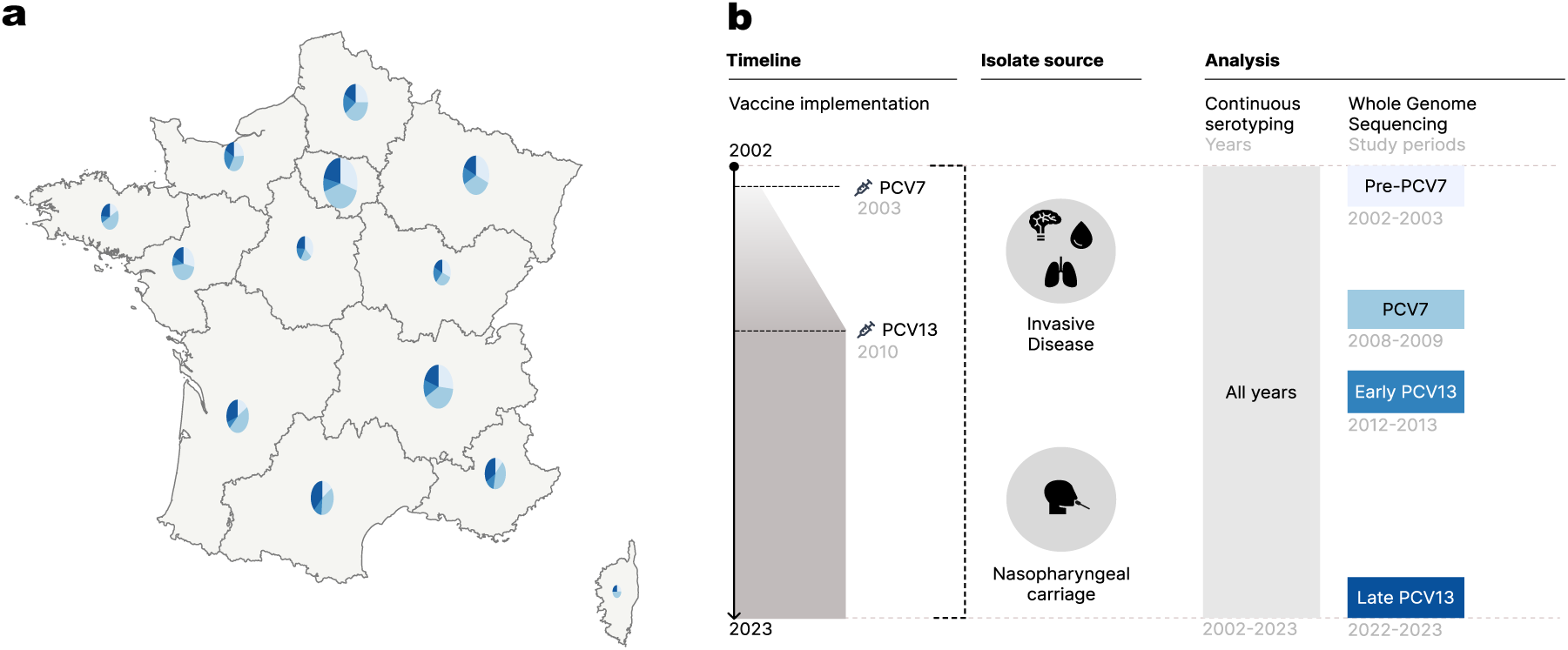
Study design. **a**, Geographic distribution of isolates included in the genomic study. Pie charts indicate the distribution of cases across study periods for each French region. The area of each pie chart is proportional to the number of samples collected in the region across all periods. **b**, Grey shading indicates the progressive implementation of PCV7 from 2003 and the implementation of PCV13 in 2010 in France. Pneumococcal invasive disease and carriage isolates were collected and serotyped from 2002 to 2023, and whole genome sequencing was performed on isolates collected during four vaccine-related periods. **a**,**b**, Study periods are represented by a gradient of blue, from lightest to darkest: pre-PCV7 (2002–2003), PCV7 period (2008–2009), early PCV13 period (2012–2013), and late PCV13 period (2022–2023). IPD, invasive pneumococcal disease; PCV, pneumococcal conjugate vaccine.

#### Genomic population structure

From this collection, we whole genome sequenced all pneumococcal carriage and IPD isolates collected during four periods reflecting successive vaccine eras between 2002 and 2023: pre-PCV7 (2002–2003), PCV7 (2008–2009), early PCV13 (2012–2013) and late PCV13 (2022–2023). 2401 genomes were included in the analysis, comprising 1397 carriage isolates and 1004 invasive isolates (**Fig. 1** and Supplementary Results 1).

Among children with pneumococcal carriage (median age = 1.1 year; interquartile range (IQR) = 0.79–1.4), PCV7 uptake was 99.2% during the PCV7 period, and PCV13 uptake was ≥99% during both the early and late PCV13 periods, consistent with the high vaccine coverage reported in France^36^. Among children with IPD (median age = 1.5 years; IQR = 0.67–5.3), 304 (30.3%) isolates were obtained from cerebrospinal fluid and 699 (69.6%) from blood. Clinical and microbiological characteristics of children included in the genomic analysis are provided in Supplementary Results 1.

De novo genome assemblies ranged from 1.95 to 2.25 Mb. Read and assembly metrics indicated high sequencing quality (Supplementary Results 2). Population clustering analysis identified 95 GPSCs, corresponding to genetically defined pneumococcal lineages. Core-genome phylogenetic analysis, performed with R packages *ape* and *phangorn*^37,38^, showed that GPSCs are largely monophyletic, meaning that each lineage corresponds to a clade of the phylogenetic tree. Twenty-three high-frequency GPSCs, each representing more than 1% of the isolates, together accounted for 82.6% of the total collection. The most prevalent lineages were GPSC10 (13.5%), GPSC6 (7.2%), GPSC7 (5.7%), and GPSC9 (5.2%). Some GPSCs were mainly observed among carriage isolates (GPSC7, GPSC59, GPSC99) whereas others were predominantly detected in IPD (GPSC15, GPSC31). Several GPSCs were associated with a single serotype (for example, GPSC31–1, GPSC12–3, GPSC15–7F, GPSC35–10A, GPSC99–21, and GPSC19–22F), whereas the majority encompassed multiple serotypes (**Fig. 2** and Supplementary Results 3). The serotype composition of high-frequency lineages appeared to shift across study periods, from vaccine to non-vaccine serotypes (**Fig. 3**)

**Fig 2.**
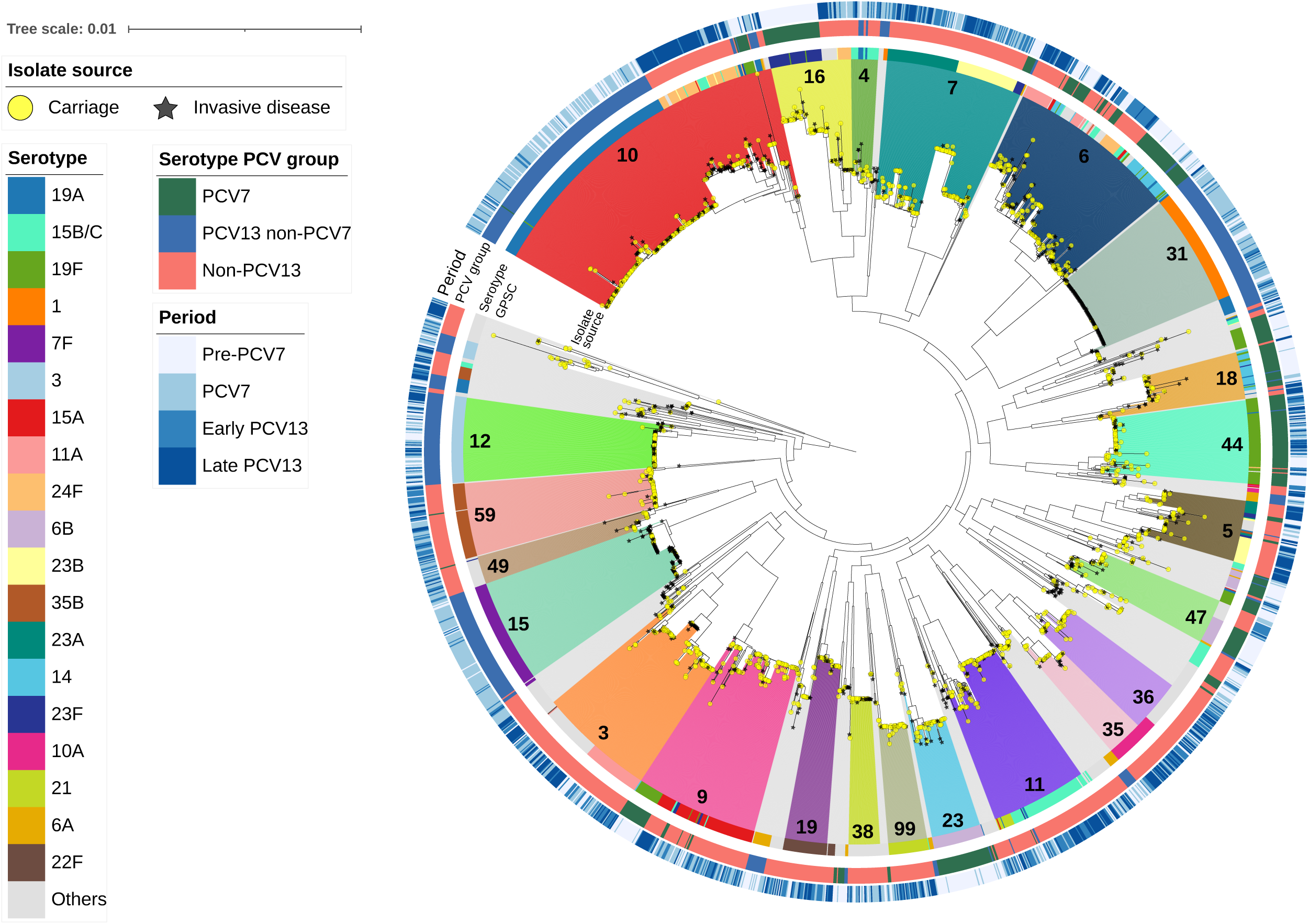
Descriptive summary of *S. pneumoniae* isolates. Midpoint-rooted maximum-likelihood phylogeny inferred from a core genome alignment. The phylogeny comprises 2401 isolates collected from the nasopharynx of children aged ≤2 years (yellow dots, n = 1397) and from sterile sites of children aged ≤17 years with invasive pneumococcal disease (black stars, n = 1004) across four vaccine-related periods in France. GPSCs representing >1% of isolates are individually annotated and visualised as coloured clade ranges, whereas low-frequency GPSCs are shown in grey. Inner-to-outer-coloured strips indicate serotype (only serotypes representing >2% of isolates are coloured and shown in the legend), serotype PCV group, and vaccine-related period. GPSCs corresponded to monophyletic groups on the phylogeny. Rare exceptions of apparent non-monophyly are discussed in Supplementary Results 3. GPSC, Global Pneumococcal Sequence Cluster; PCV, pneumococcal conjugate vaccine.

**Fig 3.**
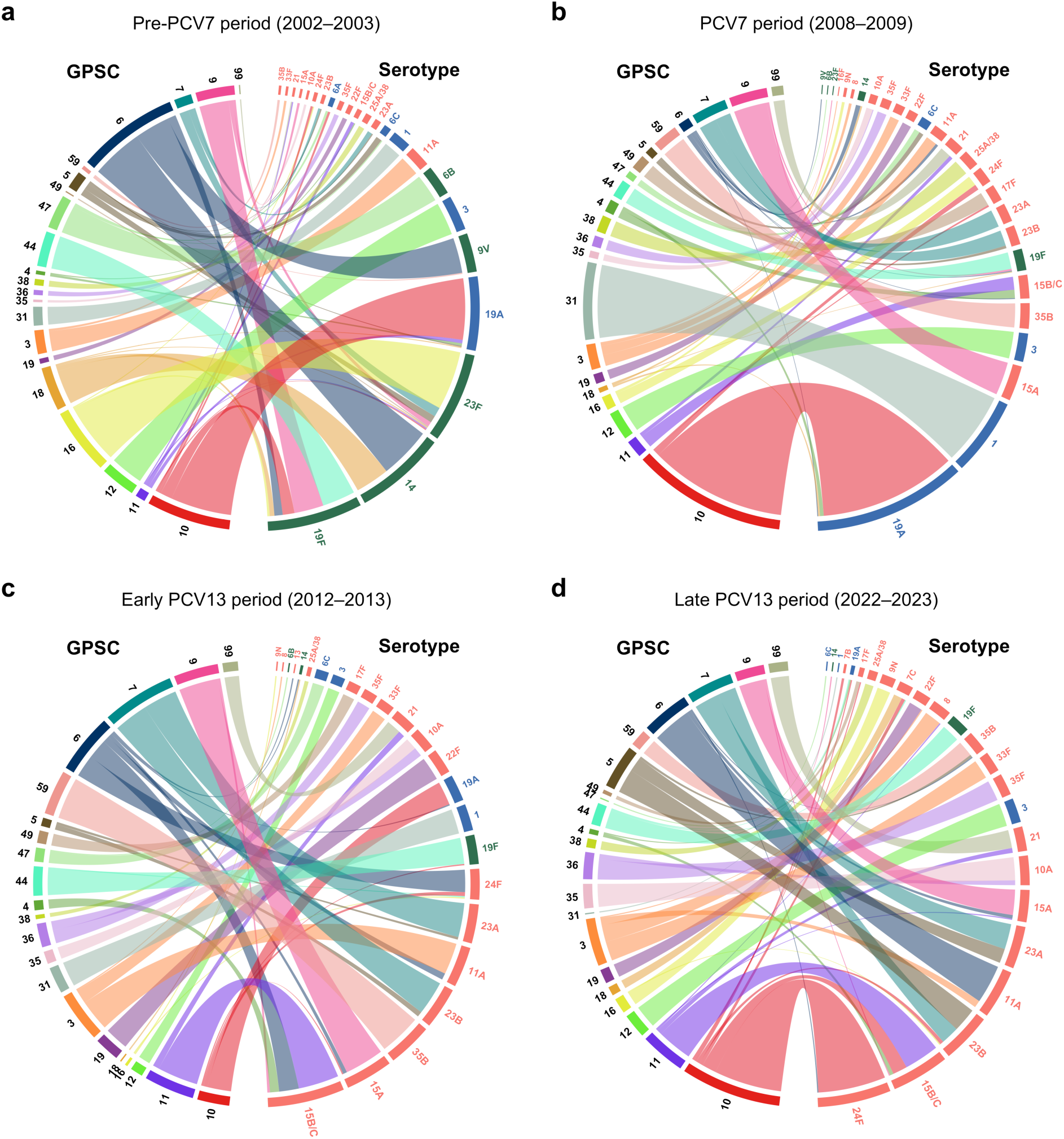
Serotype composition of persistent pneumococcal lineages in carriage and invasive disease isolates (n = 2401). **a–d,** Chord diagrams show the distribution of pneumococcal serotypes within GPSCs across the study periods: pre-PCV7 (2002–2003, **a**), PCV7 (2008–2009, **b**), early PCV13 (2012–2013, **c**), and late PCV13 (2022–2023, **d**). GPSCs are displayed in the left half of each circle; serotypes are shown in the right half and are ordered by increasing frequency. Ribbons connect serotypes to the GPSCs in which they were observed, with widths proportional to the number of isolates. Only high-frequency GPSCs (>1% of isolates) detected in all four periods are shown. Serotype labels are coloured according to vaccine group: PCV7 serotypes (green), PCV13 non-PCV7 serotypes (blue), and non-PCV13 serotypes (red). PCV, pneumococcal conjugate vaccine; GPSC, Global Pneumococcal Sequence Cluster.

### Serotype replacement in carriage and invasive disease

We leveraged two decades of carriage and IPD surveillance data covering pre- and post-PCV7 and PCV13 periods in the paediatric population to quantify serotype replacement and model the long-term temporal dynamics of vaccine and non-vaccine serotypes. We fitted temporal trends in serotype carriage and IPD rates between 2002 and 2023 using polynomial logistic regressions. Model coefficients and diagnostics are detailed in Supplementary Results 4.

Those dynamics reflected serotype replacement: PCV7 serotypes declined dramatically over time, non-PCV13 serotypes increased progressively toward a plateau, and PCV13 non-PCV7 serotypes displayed a bell-shaped trajectory combining a pre-PCV13 increase and a post-PCV13 decrease. Carriage and IPD dynamics were broadly concordant, with discrepancies in the magnitude of changes. The expansion of non-PCV13 serotypes was less prominent and the temporary increase in PCV13 non-PCV7 serotypes was more pronounced in IPD than in carriage (**Fig. 4**). The estimated total IPD incidence in 2023 (2.86 per 100,000 children aged ≤17 years; 95% confidence interval (CI) = 2.6–3.2) was strongly reduced compared with 2002 (4.8 per 100,000; 95% CI = 4.4–5.2), while carriage rate remained relatively stable. Overall, these findings highlight the incomplete replacement of serotypes in IPD compared to carriage.

**Fig 4.**
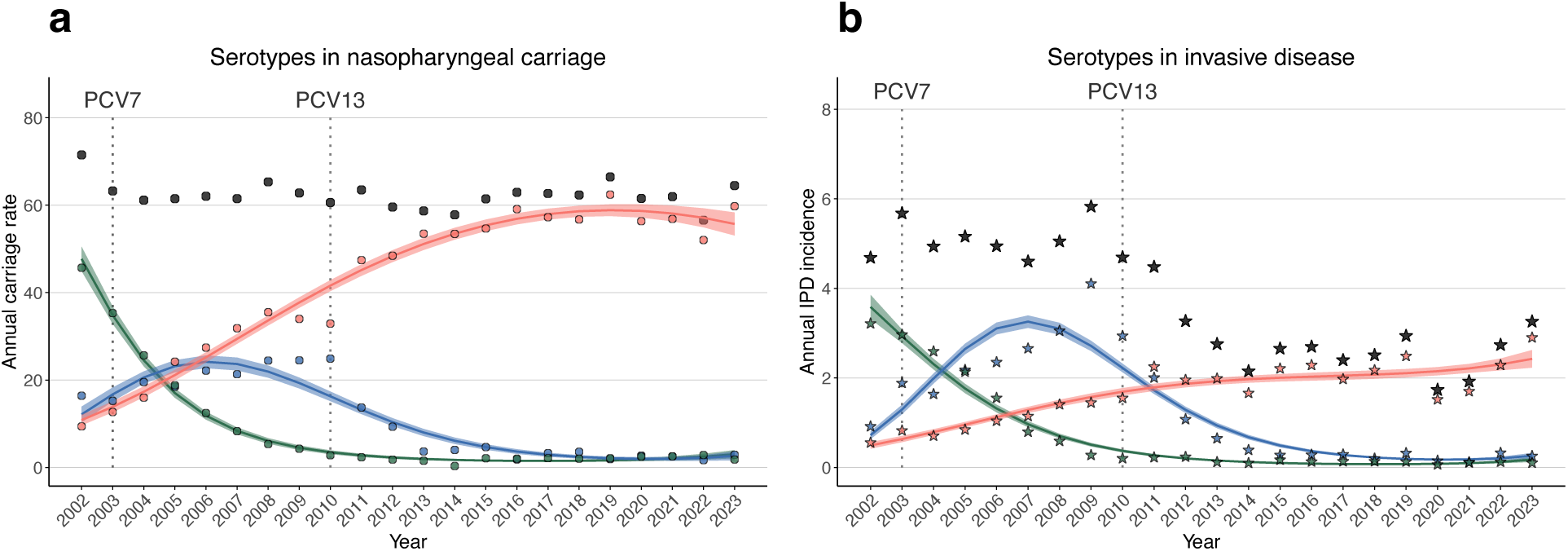
Vaccine induced serotype replacement in carriage and invasive disease. **a,b**, Observed annual carriage rate (dots) per 100 sampled children aged ≤2 years (**a**) and annual incidence of IPD (stars) per 100,000 children aged ≤17 years living in France (**b**), and model fits (curves) with 95% CIs (shaded bands) for PCV7 serotypes (green), PCV13 non-PCV7 serotypes (blue), non-PCV13 serotypes (red), and overall serotypes (black) from 2002 to 2023. Vertical dotted lines indicate the progressive implementation of PCV7 from 2003 and the introduction of PCV13 in 2010 in France. PCV, pneumococcal conjugate vaccine; IPD, invasive pneumococcal disease; CIs, confidence intervals.

### Observed evolution of pneumococcal lineages in carriage

To describe lineage dynamics depending on their pre-PCV7 (2002–2003) serotype composition, we classified lineages into three groups: PCV7-type GPSCs (>50% PCV7 serotypes), PCV13 non-PCV7-type GPSCs (>50% PCV13 non-PCV7 serotypes), and non-PCV13-type GPSCs (others), as previously published (Supplementary Table S7)^23^.

#### Between-lineage dynamics

We fitted GPSC dynamics using polynomial logistic regression on the carriage rates, including a polynomial in time and adjustment for mean GPSC penicillin minimum inhibitory concentrations (MICs), which varied between GPSCs and across periods (Supplementary Results 5). In contrast to serotypes, pneumococcal lineages remained largely stable across vaccine eras. On average, the carriage of PCV7-type GPSCs initially declined before stabilizing well above zero; that of non-PCV13-type GPSCs increased slightly before reaching a plateau; and that of PCV13 non-PCV7-type GPSCs initially increased before declining towards zero, with a slight re-increase in the late PCV13 period (**Fig. 5a–c**).

**Fig 5.**
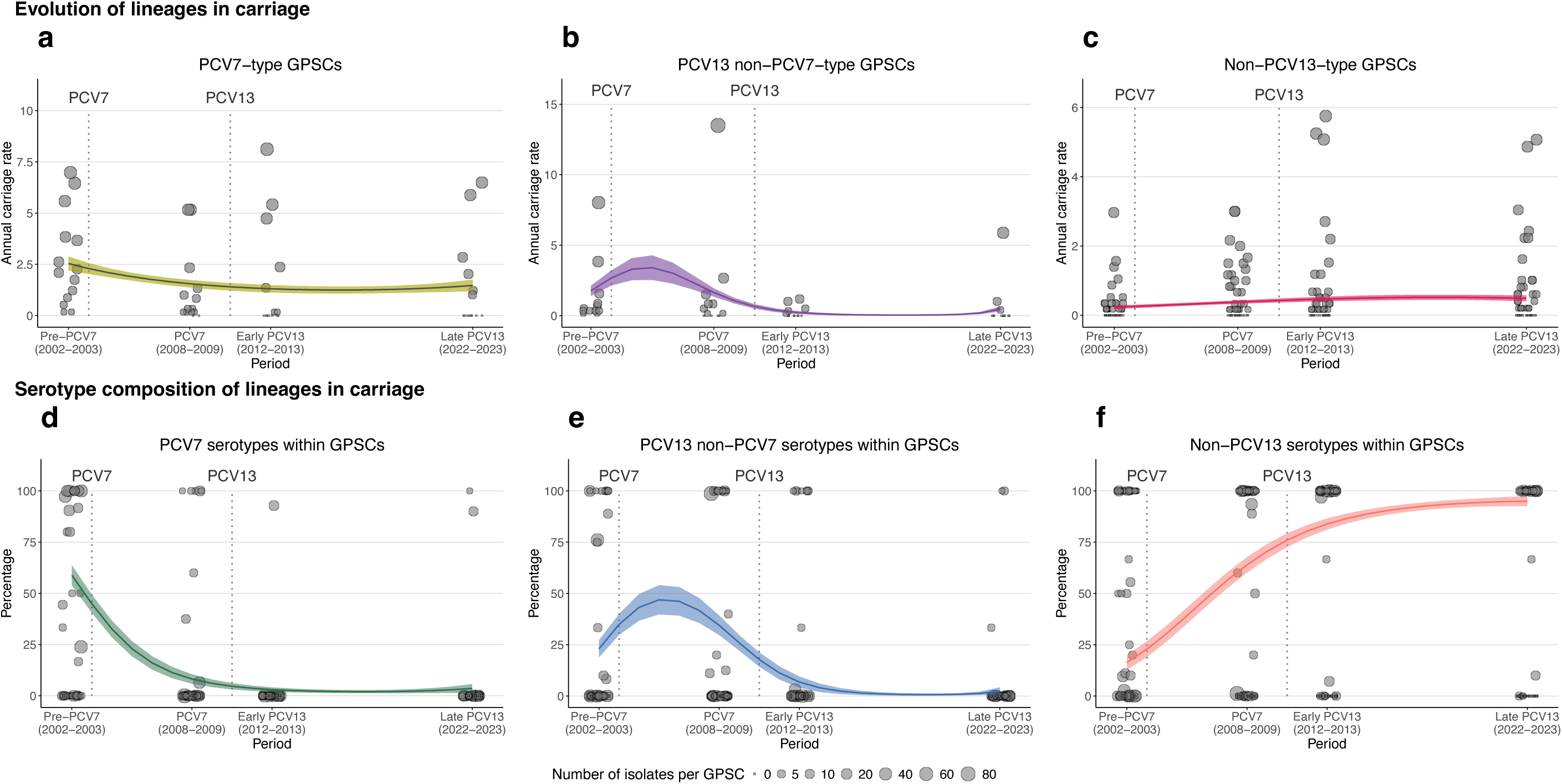
Observed evolution of pneumococcal lineages in nasopharyngeal carriage. **a–c**, Observed (dots) and model fits (curves) with 95% CIs (shaded bands) for the annual carriage rate of PCV7-type GPSCs (yellow, **a**), PCV13 non-PCV7-type GPSCs (purple, **b**), and non-PCV13-type GPCSs (pink, **c**) per 100 sampled children aged ≤2 years across vaccine eras (2002–2023). GPSCs were classified according to their pre-PCV7 (2002–2003) serotype composition (>50%). The scale of the y axes is different between the three plots. **d–f**, Observed (dots) and model fits (curves) with 95% CI (shaded bands) for the proportion of PCV7 serotypes (green, **d**), PCV13 non-PCV7 serotypes (blue, **e**), and non-PCV13 serotypes (red, **f**) within GPSCs in nasopharyngeal carriage over time (2002–2023). **a,f**, Each dot represents one GPSC, and dot size is proportional to the number of isolates per GPSC in each period. Vertical dotted lines indicate the progressive implementation of PCV7 from 2003 and the introduction of PCV13 in 2010 in France. PCV, pneumococcal conjugate vaccine; GPSC, Global Pneumococcal Sequence Cluster; CIs, confidence intervals.

#### Within-lineage serotype composition

To investigate within-lineage dynamics, we computed the relative proportions of PCV7, PCV13 non-PCV7, and non-PCV13 serotypes within each GPSC lineage across the study periods. Then, we analysed the variation of these proportions over time using polynomial logistic regressions (Supplementary Results 5). Within-lineage dynamics showed similar temporal patterns as overall serotype carriage: PCV7 serotypes dramatically declined, while non-PCV13 serotypes rose towards being hegemonic in many lineages, and PCV13 non-PCV7 serotypes displayed a similar initial increase before declining as well (**Fig. 5d–f**).

### Simulation of counterfactual scenarios

We hypothesized two processes underlying the shift observed in *S. pneumoniae* serotype composition: the replacement of serotypes within persistent lineages or the replacement of the lineages themselves. To quantify the relative contribution of these two processes to the observed lineage dynamics in carriage, we constructed two counterfactual scenarios in which one level of population structure was held constant over time. Under the “serotype-fixed” scenario, the distributions of PCV7, PCV13 non-PCV7, and non-PCV13 serotype groups within each GPSC remained constant at their pre-PCV7 (2002–2003) level. The counterfactual GPSC carriage rates then varied accordingly, so that the population-level serotype-group distribution over time matched that of the data. By contrast, under the “lineage-fixed” scenario, GPSC carriage rates remained constant at their pre-PCV7 level.

#### Serotype-fixed scenario

Counterfactual GPSC group trajectories were estimated using polynomial logistic regression models adjusted for mean GPSC penicillin MIC. Under this scenario, GPSC dynamics closely mirrored serotype-level trends: PCV7-type GPSCs declined sharply, non-PCV13-type GPSCs increased progressively toward a plateau, and PCV13 non-PCV7-type GPSCs displayed a bell-shaped trajectory (**Fig. 6a–c**). However, these inferred trajectories deviated substantially from data-informed estimates of GPSC dynamics (**Fig. 7a–c** and Supplementary Results 6).

**Fig 6.**
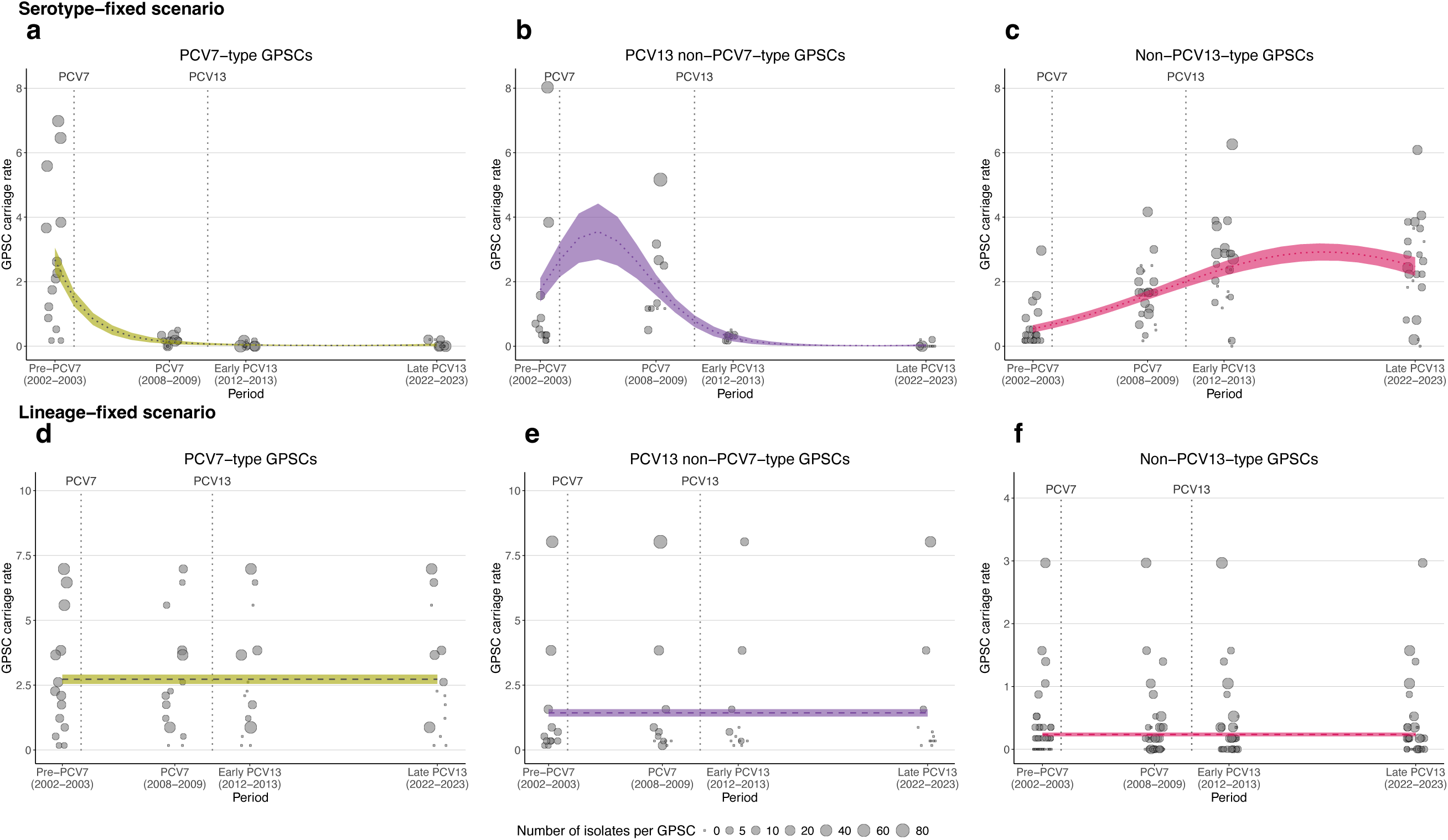
Counterfactual evolution of pneumococcal lineages in nasopharyngeal carriage. **a–f**, Counterfactual annual GPSC carriage rates (per 100 sampled children aged ≤2 years) across vaccine eras (2002–2023), together with model estimates inferred under the serotype-fixed scenario (dotted curves, **a–c**), in which within-GPSC serotype composition was held constant over time, and under the lineage-fixed scenario (dashed curves, **d–f**), in which GPSC carriage rates were fixed across periods. Shaded bands represent 95% CIs. GPSCs were grouped according to their pre-PCV7 (2002–2003) serotype composition (>50%) as PCV7-type (yellow, **a,d**), PCV13 non-PCV7-type (purple, **b,e**), and non-PCV13-type (pink, **c,f**). Each dot represents one GPSC, with dot size proportional to the observed number of isolates in each period; all GPSCs are shown in grey. Vertical dashed lines indicate the progressive implementation of PCV7 from 2003 and the introduction of PCV13 in 2010. PCV, pneumococcal conjugate vaccine; GPSC, Global Pneumococcal Sequence Cluster; CIs, confidence intervals.

**Fig 7.**
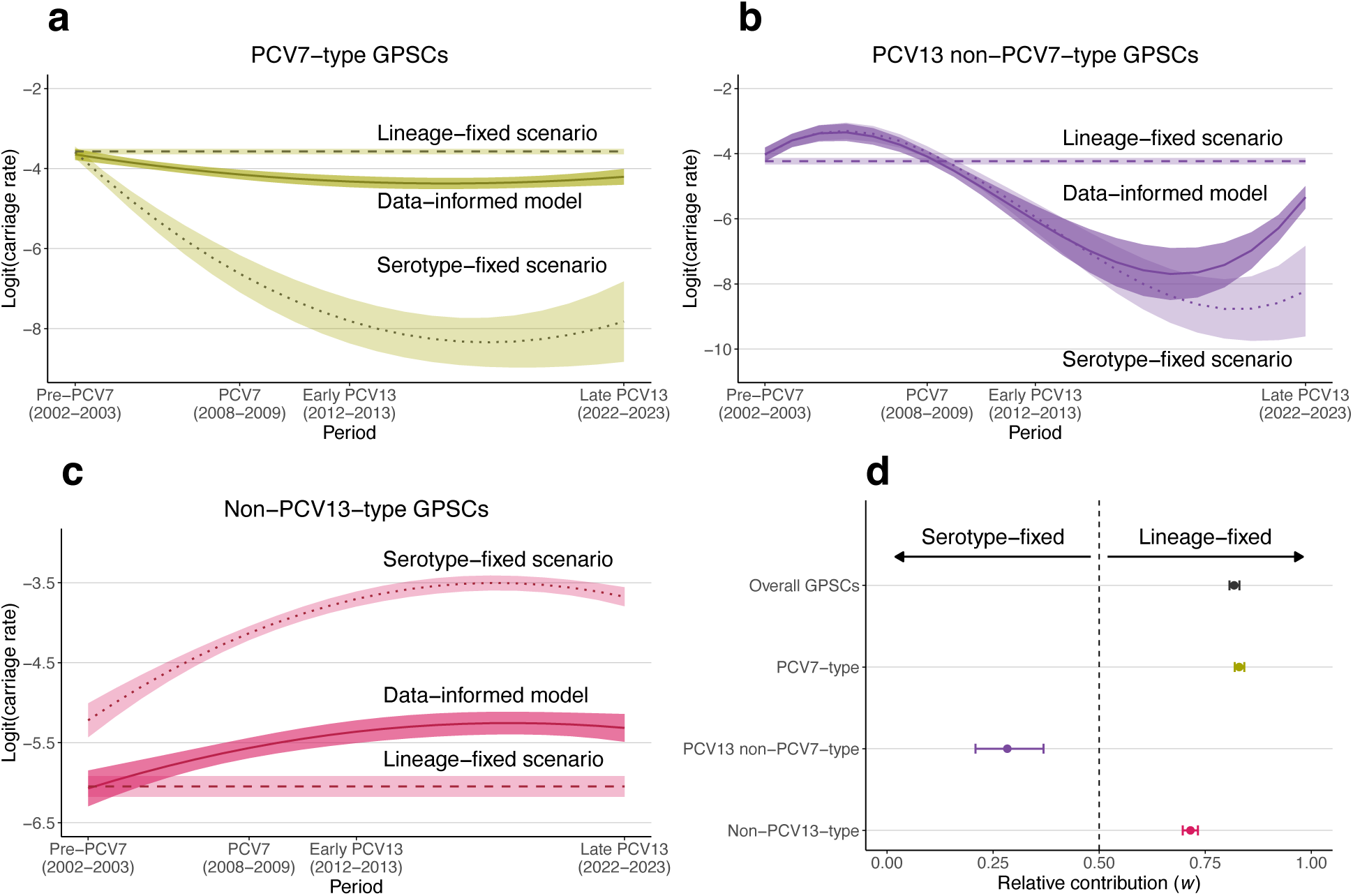
Modelled carriage dynamics of lineages under mechanistic scenarios. **a–c**, Data-informed model estimates (solid curves) based on observed GPSC carriage rates, compared with the serotype-fixed scenario (dotted curves), in which within-GPSC serotype composition is held constant over time, and the lineage-fixed scenario (dashed curves), in which GPSC carriage rates are fixed across periods. Trajectories are shown for PCV7-type GPSCs (yellow, **a**), PCV13 non-PCV7-type GPSCs (purple, **b**), and non-PCV13-type GPSCs (pink, **c**) from 2002 to 2023. Shaded bands indicate 95% CI. Y axes show logit-transformed annual carriage rates per 100 sampled children aged ≤2 years, consistent with the scale used for contribution calculations. **d**, Relative contribution *w*_G_ of each scenario to the data-informed GPSC dynamics, estimated by minimizing the weighted sum of squared deviations between the combination of scenario trajectories and the data-informed model.

#### Lineage-fixed scenario

Under this scenario, inferred GPSC carriage rates were held constant across study periods. GPSC group trajectories were estimated using polynomial logistic regressions adjusted for mean GPSC penicillin MIC. Except for PCV13 non-PCV7-type GPSCs, estimated GPSC carriage trends were broadly consistent with those derived from the data-informed model, (**Fig. 6d–f**, **Fig. 7a–c**, and Supplementary Results 6).

#### Quantifying mechanism contributions

To quantify the relative contributions of lineage replacement and within-lineage serotype replacement, we approximated observed lineage dynamics as a weighted combination of the lineage-fixed and serotype-fixed scenarios. The weighted parameter *w*_G_ varied between 0 (only takes the serotype-fixed scenario into account) and 1 (only takes the lineage-fixed into account). We estimated the value of *w*_G_ minimizing the squared deviation between the weighted combination of scenarios and the data-informed trajectory across 2002–2023.

The contribution of the lineage-fixed mechanism was high for both PCV7-type GPSCs (*w_PCV7_* = 0.83, 95% CI = 0.82–0.84) and non-PCV13-type GPSCs (*w*_non- *PCV13*_ = 0.72, 95% CI = 0.70–0.73). This contribution was markedly lower for PCV13 non-PCV7-type GPSC dynamics (*w_PCV13_* = 0.28, 95% CI = 0.21–0.37). Overall, when considering all three GPSC groups, the contribution of the lineage-fixed scenario to the data-informed lineage dynamics was *w*_’*overall*_ = 0.82 (95% CI = 0.81–0.83), whereas that of the serotype-fixed scenario was 1-*w_overall_* = 0.18 (**Fig. 7d**).

### Predicting serotypes in invasive disease

IPD incidence corresponds to the product of carriage rate by serotype-specific invasive disease potential^7^. To investigate the consequences of each mechanism on serotype evolution in IPD, we estimated IPD incidence based on each scenario carriage prediction and serotype-specific invasive disease potentials retrieved from the literature^7–10^.

Focusing on the late PCV13 period (2022–2023), the IPD estimates from the lineage-fixed scenario were strongly correlated with observed serotype-level IPD incidence (Pearson’s r = 0.82, 95% CI = 0.70–0.89; *P* < 0.001), whereas the estimates from the serotype-fixed scenario showed no significant correlation (Pearson’s r = 0.09, 95% CI = -0.19–0.36; *P* = 0.518) (**Fig. 8**).

**Fig 8.**
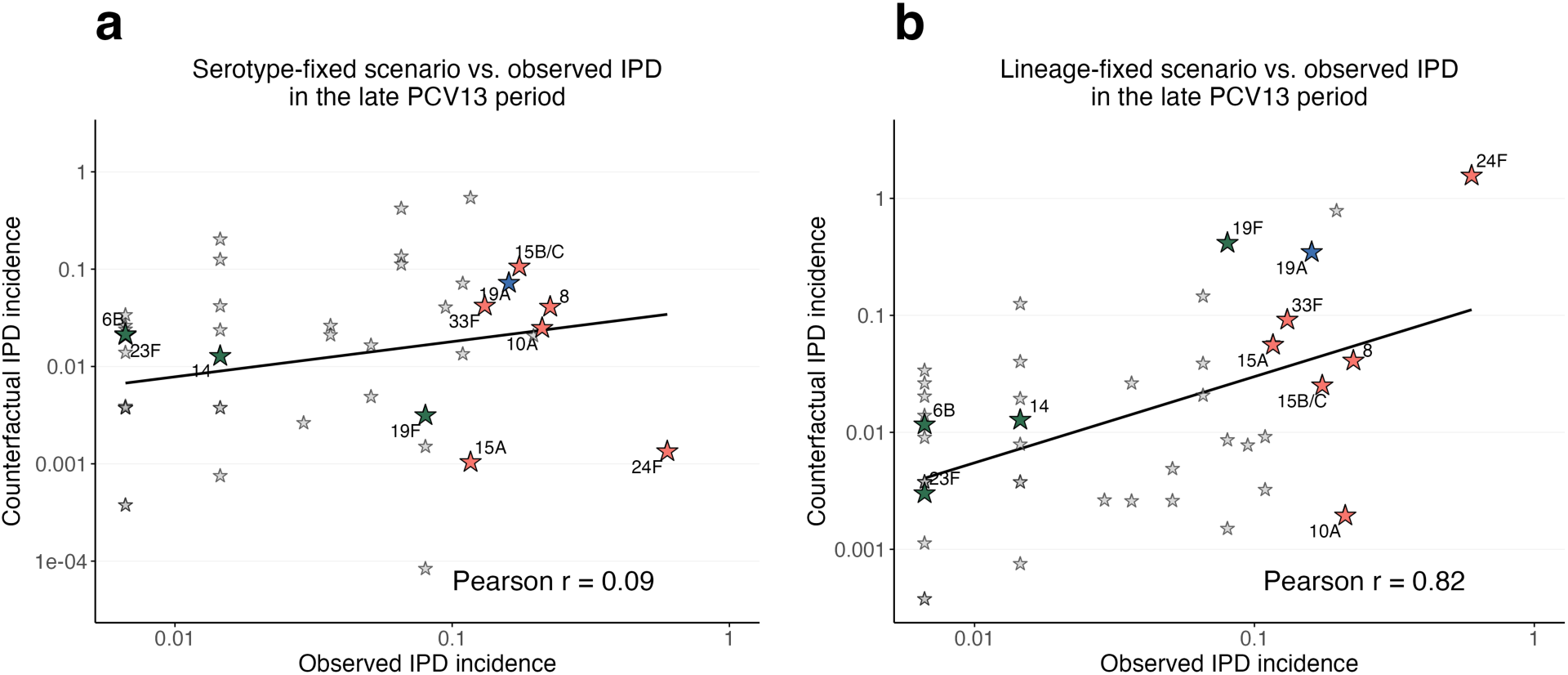
Correlation between observed and estimated serotype-specific IPD incidence from carriage under counterfactual scenarios. **a**,**b**, Correlation between observed IPD incidence and IPD incidence estimated from carriage under the serotype-fixed scenario (**a**) and the lineage-fixed scenario (**b**) during the late PCV13 period (2022–2023). Each point represents a serotype. Selected serotypes were annotated, including those most frequent before PCV7 introduction and those emerging in the late PCV13 period. Colours indicate PCV7 (green), PCV13 non-PCV7 (blue), and non-PCV13 (red) serotypes. Solid lines show linear regression fits. Pearson correlation coefficients are indicated. Y axes are in log scale. IPD, invasive pneumococcal disease; PCV, pneumococcal conjugate vaccine.

We then examined the estimated IPD incidence across all periods. Among the most frequent serotypes in the pre-PCV7 period (14, 6B, 19A, 19F, 23F, and 9V), both scenarios produced estimates close to the observed serotype-specific trends in IPD. By contrast, when focusing on the main emerging serotypes in the late PCV13 period (24F, 15B/C, 10A, 33F, 8, and 15A), the lineage-fixed scenario more accurately captured the rise of serotype 24F, which has become a major cause of IPD. Other emerging serotypes were captured by one or the other scenario (**Fig. 9**).

**Fig 9.**
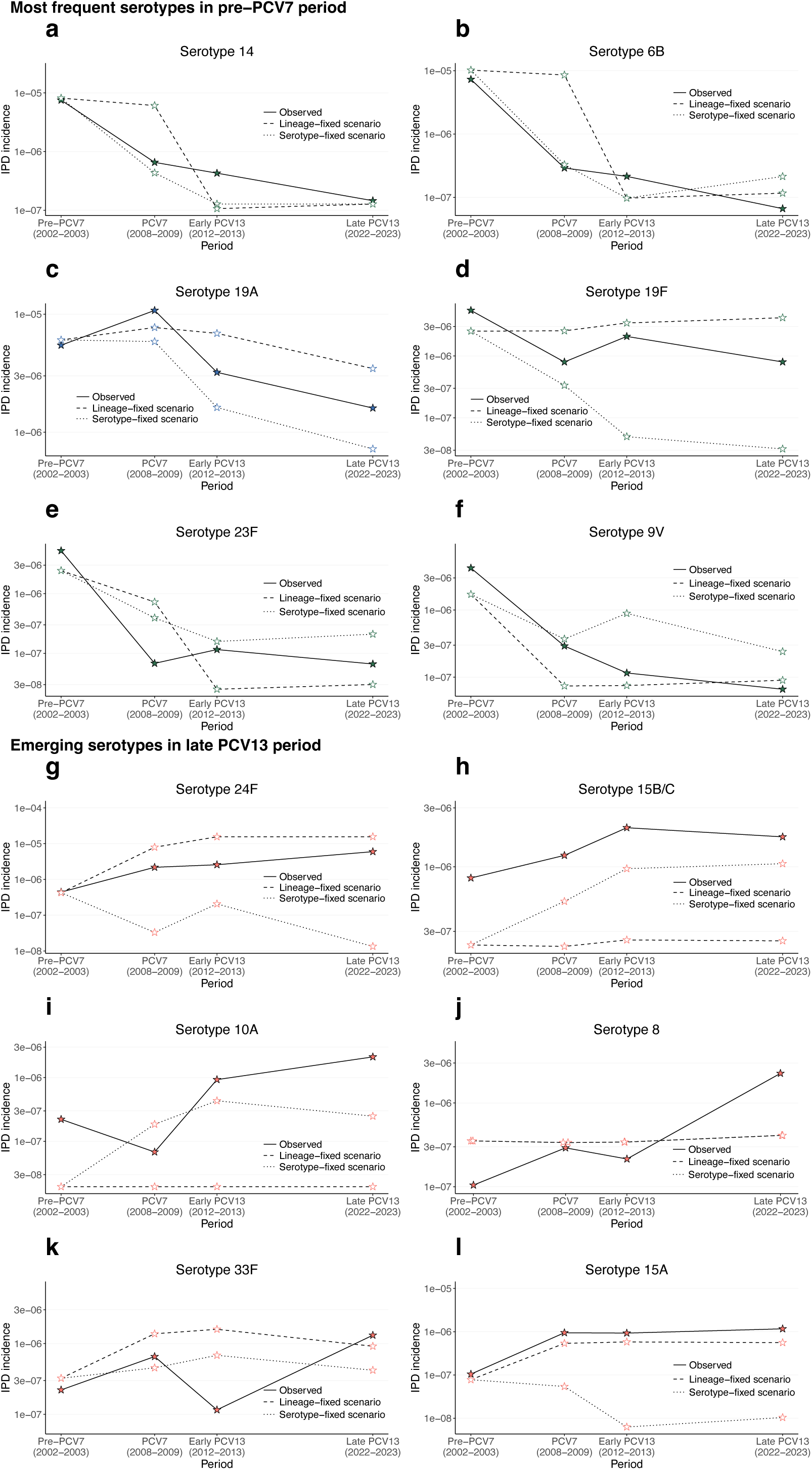
Observed and estimated IPD incidence of serotypes under counterfactual scenarios. **a–l**, Observed IPD incidence (filled stars and solid lines) and incidence estimated from carriage (open stars) across study periods under the serotype-fixed (dotted lines) and lineage-fixed (dashed lines) scenarios for the most frequent serotypes in the pre-PCV7 period (**a–f**) and the main emerging serotypes in the late PCV13 period (**g–l**). Colours indicate PCV7 (green), PCV13 non-PCV7 (blue), and non-PCV13 (red) serotypes. Incidence is expressed as the number of IPD cases per 100,000 children aged ≤17 years. Y axes are in log scale. **j**, Lineage-fixed and serotype-fixed scenarios are confounded for serotype 8. IPD, invasive pneumococcal disease; PCV, pneumococcal conjugate vaccine.

## Discussion

In this study, we investigated the relative contributions of lineage turnover and serotype replacement within persistent lineages to *S. pneumoniae* serotype replacement following the introduction of PCV7 and PCV13. Despite pronounced changes in their serotype composition, pneumococcal lineages remained remarkably stable in carriage over two decades. Carriage dynamics were more consistent with the lineage-fixed scenario than with the serotype-fixed scenario. These findings indicate that changes in serotype distribution occurred primarily through serotype replacement within relatively stable genetic lineages rather than through widespread lineage turnover. This mechanism strongly shaped the emergence of predominant non-vaccine serotypes, including serotype 24F which expanded within the multidrug-resistant and virulent GPSC10 and is now among the most frequent serotypes causing IPD in several countries^19,39^.

By jointly analysing nasopharyngeal carriage and IPD, we provide a more complete view of lineage and serotype dynamics, building on previous studies reporting the expansion of non-vaccine serotypes within vaccine-type GPSCs^23,25^. For example, in South Africa, GPSC5 was mainly associated with vaccine serotype 23F before PCV13 and persisted across vaccine periods through expansion of the non-vaccine serotype 35B/D, while instances of “clonal replacement” have also been reported^23^. However, analyses relying solely on IPD data provide only a partial view of pneumococcal population dynamics, as disease surveillance is influenced by the invasive disease potential of individual serotypes^26–29^. In our dataset, GPSC5–23F was primarily replaced in carriage by GPSC5–23A and GPSC5–23B, whereas GPSC5–35B/D was observed in IPD. This joint analysis of carriage and IPD therefore enables us to overcome this observational filter and to quantify the mechanisms underlying serotype replacement.

The relative contribution of these mechanisms may vary with epidemiological and vaccination implementation settings. In contrast to the overall stability observed across most GPSCs, dynamics among PCV13 non-PCV7-type GPSCs showed greater deviation from the lineage-fixed scenario, suggesting greater variation in lineage carriage rate over time. This pattern was largely driven by GPSC10 (163/275, 59.3% of PCV13 non-PCV7 isolates over the whole period), which exhibited substantial fluctuations in carriage rate simultaneously to shifts in serotype composition. Initially dominated by serotypes 19A and 19F, GPSC10 expanded during the PCV7 period as 19A increased, declined rapidly following PCV13 introduction in parallel with reductions in 19A, and later resurged with non-PCV13 serotype 24F. These dynamics occurred in the context of rapid nationwide implementation of PCV13 in France, which may have imposed immediate selective pressure, limiting the time available for lineages to adjust their serotype composition. By contrast, the more gradual implementation of PCV7 between 2003 and 2007 may have allowed PCV7-type lineages to persist through progressive shifts towards non-vaccine serotypes. These observations suggest that the balance between lineage turnover and within-lineage serotype replacement may depend on the regional epidemiology, as well as on timing, speed and extent of PCV implementation.

Our data indicate that vaccine-driven serotype replacement primarily reflects changes in the frequency of serotypes within existing pneumococcal lineages, rather than the emergence of new variants through capsular switching. The detection of lineage-serotype combinations, such as GPSC5–23A, GPSC5–23B, and GPSC10–24F, prior to PCV introduction supports the expansion of variants already present at low frequency, whereas capsular switching represents, in most cases, historical events predating vaccine introduction^25^. This pattern is consistent with the action of negative frequency-dependant selection, which has been suggested to maintain stable combinations of accessory genes within pneumococcal lineages^40^. Vaccination thus acted as a selective filter on pre-existing genetic diversity, reshaping serotype composition within lineages while preserving the genomic structure of pneumococcal populations.

Classifying GPSCs based on their serotype composition does not provide a stable representation over time. By deliberately classifying GPSCs according to their pre-PCV7 serotype composition, as previously described^23^, we assessed whether vaccine serotype categories align with lineage structure. This alignment proved unstable: lineages initially defined as PCV7-type frequently shifted towards predominance of non-PCV7 serotypes under vaccine pressure. For example, GPSC6 persisted through progressive replacement of vaccine serotypes (9V, 14, 19F, and 19A) by non-PCV13 serotypes (11A, 15A, and 35B). These findings indicate that serotype composition alone does not capture intrinsic lineage structure and suggest that the diversity of serotypes within lineages may better reflect their evolutionary potential under vaccine pressure. Rather than focusing on dominant serotypes, a quantitative assessment of serotype diversity expressed within each lineage could provide a more informative measure of its capacity to adapt, with more diverse lineages being more likely to persist through shifts in serotype composition. Predicting pneumococcal evolution under vaccine pressure may therefore require grouping lineages according to both their serotype diversity and their ecological success in carriage, thereby identifying those most likely to persist and generate emerging serotypes.

Similar patterns of lineage persistence despite shifts in capsular type have been described in other bacterial species. *N. meningitidis*, another nasopharyngeal commensal capable of invasive disease, illustrates this phenomenon: lineage 11 has been associated with serogroups B, C, W and Y in different epidemiological settings, showing how stable clonal backgrounds can express multiple serogroups over time and persist under changing immune pressures^5^. In contrast, *H. influenzae*, which also occupies the child’s nasopharynx, exhibits a population structure akin to panmixia, with extensive recombination limiting deep genetic divergence and the formation of stable lineages. This may explain why no distinct lineage expanded following the decline of *H. influenzae* b after conjugate vaccination^4^. Although both *H. influenzae* and *S. pneumoniae* are highly recombinogenic^41^, their population structures differ fundamentally. Pneumococci are organized into stable, deeply divergent lineages in which historical recombination has generated capsular diversity^42,43^. Together, these observations suggest that the capacity of bacterial populations to adapt to vaccination is shaped by their population structure, with pneumococci favouring shifts in serotype composition within stable lineages.

It should be noted that our counterfactual scenarios rely on simplifying assumptions that do not fully capture biological complexity. In both scenarios, serotypes not observed within a lineage pre-PCV7 could not emerge, thus leading to discrepancies in the emergence of non-vaccine serotypes compared to reality. In the serotype-fixed scenario, the approach based on serotype groups (PCV7, PCV13 non-PCV7 and non-PCV13) rather than individual serotypes lead to the emergence of non-vaccine serotypes different from the actual ones. In the lineage-fixed scenario, actual emerging serotypes did emerge, but only in lineages where they were initially observed. Thus, our scenarios did not account for potential lineage–serotype-specific interactions influencing carriage fitness.

Furthermore, carriage data were obtained from children aged ≤2 years, whereas IPD data included children up to 17 years of age. Although this age mismatch may introduce additional uncertainty when translating carriage dynamics into disease outcomes, carriage in young children is strongly associated with IPD patterns in other age groups^44^.

Current PCV strategies rely on the progressive expansion of serotype valency to reduce the burden of IPD. With higher-valency formulations such as PCV15 and PCV20 now being introduced into paediatric immunization programmes, anticipating their epidemiological impact has become increasingly important. To date, most studies aiming to predict the impact of new PCVs have focused on vaccine serotype coverage in relation to serotype distribution in IPD^18,45,46^. However, vaccine-driven changes in pneumococcal populations do not necessarily eliminate successful lineages, which may instead persist and expand through alternative serotypes. The emergence of GPSC10–serotype 24F in France illustrates this process, as this lineage–serotype combination proved highly successful in both carriage and invasive disease and was associated with elevated antimicrobial resistance^19^. Our findings suggest that anticipating the impact of next- generation PCVs will require moving beyond serotype-level approaches to incorporate lineage-level dynamics.

Pneumococcal populations respond to vaccine pressure primarily through shifts in serotype composition within stable genetic lineages. Our findings provide a quantitative framework for understanding pathogen adaptation under vaccine selection and highlight the importance of integrating lineage structure and carriage ecology to improve predictions of serotype emergence following the introduction of higher-valency pneumococcal vaccines.

## Methods

### Ethical approval

The pneumococcal carriage surveillance study was approved by the Saint-Germain-en-Laye Hospital Ethics Committee and registered at ClinicalTrials.gov (NCT04460313). Written informed consent was obtained from parents or legal guardians of participating children. The IPD surveillance study was approved by the French National Data Protection Commission (approval reference number 999298). As this study was part of ongoing national surveillance coordinated by the French National Public Health Agency, it did not require additional ethical committee approval or written informed consent.

### Study design and data collection

We conducted a whole-genome sequencing study of pneumococcal isolates collected through two nationwide active surveillance networks monitoring nasopharyngeal carriage and IPD in France from 2002 to 2023, encompassing pre- and post-PCV implementation periods.

From January 2002 to December 2023, nasopharyngeal samples were collected through a prospective ambulatory surveillance network involving approximately 160 paediatricians across France. Children aged ≤2 years presenting with acute otitis media associated with fever and/or otalgia and/or irritability were eligible for inclusion, as previously described^14^.

In France, pneumococcal isolates recovered from IPD – defined as isolation from a normally sterile site – in children are referred to the NRCP for serotyping and antimicrobial susceptibility testing. The national IPD surveillance system involves more than 250 hospital laboratories and has been estimated, through capture-recapture analyses, to cover 73% to 79% of inpatient hospital stays for IPD^12^. Surveillance network information, clinical criteria, sampling procedures, and microbiological methods are detailed in Supplementary Methods 1.

Demographic and clinical data were collected in both surveillance systems and included age, sex, site and date of isolation, and antimicrobial susceptibility testing results, including MICs.

From this collection, we selected for whole-genome sequencing all pneumococcal carriage and IPD isolates collected during four predefined periods corresponding to successive vaccine eras: pre-PCV7 (2002–2003), PCV7 (2008–2009), early PCV13 (2012–2013), and late PCV13 (2022–2023) (Supplementary Methods 2).

### Whole-genome sequencing and genomic characterization

Selected isolates were whole-genome sequenced using an Illumina NovaSeq^™^ 6000 platform (Illumina^®^, San Diego, USA). We characterized each genome by assigning GPSC using PopPUNK (version 2.7.5)^47^. Serotypes were assessed *in vitro* with agglutination and *in silico* with SeroBA (version 2.0.5)^48^. Details of DNA extraction, sequencing, quality control, genome assembly and analysis are provided in Supplementary Methods 3. All raw sequencing data generated in this study have been deposited in the NCBI Sequence Read Archive (SRA) under BioProject accession number PRJNA1453032.

### Classification of serotype and GPSC groups

Serotypes were grouped into three categories based on those covered by the successive vaccines: PCV7 serotypes (4, 6B, 9V, 14, 18C, 19F, and 23F), PCV13 non-PCV7 serotypes (1, 3, 5, 6A, 7F, and 19A), and non-PCV13 serotypes (serotypes not included in PCV13, including non-typeable isolates).

GPSCs were classified into three groups based on their serotype composition in the pre-PCV7 period (2002–2003): PCV7-type GPSCs (>50% PCV7 serotypes), PCV13 non-PCV7-type GPSCs (>50% additional PCV13 serotypes), and non-PCV13-type GPSCs (others)^23^.

### Models of serotype and lineage dynamics over time

We analysed variations over time in carriage rate and IPD incidence, for serotypes and GPSC lineages. IPD incidence was estimated based on the annual number of IPD cases in children ≤17 years in France, corrected to account for the 73% to 79% coverage of the *Santé publique France* surveillance system^12,35^. We used polynomial logistic regressions, with time as the explanatory factor. In those regressions, the response variable *Y* would be of the form:

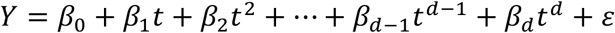

with *t* the year (centred and scaled), *d* the degree of the polynomial, *β_d_* the coefficient of *d^th^* degree term, and *ε* the residual. When considering vaccine period, we set *t* as the first year of each period (2002 for pre-PCV7, 2008 for PCV7, 2012 for early PCV13, and 2022 for late PCV13). For regressions on GPSC carriage rate, we added the log-transformed mean penicillin MIC as a covariate to account for its potential impact^23^. For serotypes, we selected the degree of the polynomial *d* (between 0 and 3 for our models) based on the Akaike information criterion^49^. We assumed that serotypes and lineages followed similar temporal patterns in carriage under vaccine selection. Therefore, we took the same degree of polynomial as that of the serotype models for the GPSC data-informed models. Additionally, we systematically performed a model diagnostic, including residual patterns and autocorrelation structures (Supplementary Methods 4).

### Construction of counterfactual scenarios

We generated two counterfactual datasets, each representing a scenario accounting for a single mechanism hypothesized to explain serotype replacement in *S. pneumoniae* carriage. In the serotype-fixed scenario, variation arose solely from changes in lineage carriage rates, whereas in the lineage-fixed scenario it arose solely from changes in serotype frequencies within lineages.

Under the serotype-fixed scenario, we fixed the relative proportion of each serotype group (PCV7, PCV13 non-PCV7 and non-PCV13) within each GPSC to their pre-PCV7 (2002–2003) values. We then numerically computed counterfactual GPSC carriage rates in each period that minimized the differences between the resulting population-level carriage rate of serotype groups and the observed ones. Under the lineage-fixed scenario, we fixed GPSC carriage rates to their pre-PCV7 levels.

We fitted the counterfactual carriage rates across periods with polynomial logistic regressions adjusted for the mean penicillin MIC of each GPSC (Supplementary Methods 5).

### Quantification of mechanism contribution

To quantify the relative contribution of the mechanisms involved in each scenario, we defined the weighted combination of the two scenarios as:

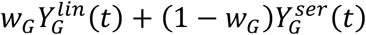

with 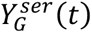 the estimated carriage of GPSCs of group *G* at year *t* under the serotype-fixed scenario, 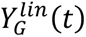 the estimated carriage under the lineage-fixed scenario, and *w*_G_ ∈ [0,1] a weight parameter reflecting the relative contribution of the lineage-fixed scenario. Hence, (1 – *w*_G_) corresponds to the relative contribution of the serotype-fixed scenario. For each GPSC group and for all GPSCs together, we numerically estimated the contribution parameter *w*_G_ minimizing the sum of squared differences between the weighted combination of the two scenarios and the estimated carriage under the data-informed model, across all years (2002–2023).

We generated the distribution of the value of *w*_!_ using parametric simulation based on the model estimates and standard errors of 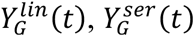 and 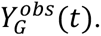 The 95% CIs corresponded to the 2.5^th^ and 97.5^th^ percentiles of this distribution (Supplementary Methods 6).

### Estimation of invasive pneumococcal disease from carriage

We estimated IPD incidence of serotype *s* in year *t* as proportional to the product of its carriage rate and its invasive disease potential:

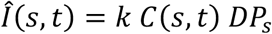

with *C*_s,t_ the carriage rate of serotype *s* in year *t*, *DP*_s_ the serotype-specific invasive disease potential, and *k* a calibration constant. For *DP*_s_, we used serotype-specific invasive disease potential estimates from previously published studies^7–10^ (Supplementary Table S3). When no estimate was available, we assigned a value of 1. We defined the calibration constant *k* such that the total estimated IPD incidence matched that of 2002.

This method predicted IPD incidence aggregated by serotype groups (PCV7, PCV13 non-PCV7 and non-PCV13) matching the observed temporal trends in IPD, thus ensuring the soundness of this carriage-based method for providing realistic approximations of observed IPD dynamics.

We used this method to estimate IPD incidence for each serotype under our two scenarios during the pre-PCV7, PCV7, early PCV13 and late PCV13 periods, based on the serotype carriage rates derived from the counterfactual GPSC carriage rates. We assessed the agreement between predicted and observed serotype-specific IPD incidence using Pearson’s correlation coefficients. Temporal trends were visually compared with observed data for the most frequent serotypes in the pre-PCV7 period and the main emerging serotypes in the late PCV13 period (Supplementary Methods 7).

All statistical analyses and visualisations were performed in R v.4.5.2^50^.

## Acknowledgements

Author guarantor statement: All authors meet the four ICMJE criteria for authorship.

## Author Contributions

ZA, TMJ, and NO designed the study. ZA, NO, RC, CL, SBe, FC, AW, SBo, PB, CR, MN, AEM, and EV were involved in the acquisition of data. ZA, TMJ, NO, and KL contributed to the data analysis. ZA drafted the manuscript. All authors made substantial contributions to the interpretation of data, provided critical revision of the manuscript for important intellectual content, and approved the final version of the manuscript. ZA, TMJ and NO take responsibility for the content of the manuscript, including the integrity of the data and the accuracy of the data analysis.

## Other contributions

We thank the children in the study and their families, all paediatricians from the PARI/ AFPA networks who participated in the study, the National Reference Centre for Pneumococci and GenoBioMICs Platform technical staffs, and *Santé publique France* for their support.

