## Supplementary Materials for "Quantifying the mechanisms of vaccine-induced serotype replacement in *Streptococcus pneumoniae* using genome-informed modelling"


***Supplementary Methods.*** **2**

***Supplementary Methods 1 | Data sources.*** *2*
*Nasopharyngeal carriage* 2
*Invasive pneumococcal disease* 2
*Microbiological investigations* 2

***Supplementary Methods 2 | Pneumococcal conjugate vaccines and study periods.*** *4*
*National immunization program* 4
*Study periods* 4

***Supplementary Methods 3 | Whole genome analysis.*** *5*

***Supplementary Methods 4 | Model selection based on AIC comparison.*** *7*

***Supplementary Methods 5 | Construction of counterfactual scenarios.*** *9*
*Numerical optimisation of GPSC carriage rates under the serotype-fixed scenario* 9

***Supplementary Methods 6 | Quantification of mechanism contribution.*** *10*

***Supplementary Methods 7 | Estimation of serotype-specific invasive pneumococcal disease from carriage scenarios.*** *11*

***Supplementary Results.*** **14**

***Supplementary Results 1 | Whole genome quality control.*** *14*
*Read quality assessment* 14
*Relationship between read-level QC and assembly quality* 14
*Genome assembly quality* 15
*Taxonomic validation* 15

***Supplementary Results 2 | Pneumococcal population structure defined by GPSCs, serotypes and core-genome phylogeny.*** *16*
*GPSC distribution* 16
*Serotype composition of GPSCs* 16
*Core-genome phylogeny and concordance with GPSC assignment* 17

***Supplementary Results 3 | Models for serotype carriage rate and IPD incidence over time.*** *19*

***Supplementary Results 4 | GPSC characteristics and evolution in carriage.*** *21*
*Penicillin non-susceptibility of GPSCs across study periods* 23
*Data-informed models for GPSC carriage rates over time* 24
*Data-informed models for within-GPSC serotype composition over time* 25

***Supplementary Results 5 | Models for counterfactual GPSC carriage rates under serotype-fixed and lineage-fixed scenarios.*** *26*

***References*** **30**

### Supplementary Methods.

#### Supplementary Methods 1 | Data sources.

*Nasopharyngeal carriage*

Between January 2001 and December 2023, pneumococcal strains were obtained from nasopharyngeal samples collected as part of a prospective, national ambulatory surveillance study conducted by the networks ACTIV (*Association Clinique et Thérapeutique Infantile du Val de Marne*, https://www.activ-france.com/en/) and AFPA (*Association Française de Pédiatrie Ambulatoire*, https://afpa.org/) since 2001[1,2].

During the winter season of each subsequent year, 160 paediatricians throughout France enrolled children aged 6 to 24 months presenting with acute otitis media (AOM) associated with fever and/or otalgia and/or irritability[1–4]. AOM was defined according to the Paradise algorithm criteria for acute suppurative otitis media (effusion plus marked redness, marked bulging or moderate redness and bulging)[5]. Nasopharyngeal carriage of *S. pneumoniae* was examined in children with AOM, as these children exhibit higher carriage rates than healthy children[6]. This approach enabled more efficient monitoring of serotype distribution and antibiotic susceptibility over time while minimizing the number of children requiring swabbing. Children who received antibiotic treatment within 7 days prior to enrolment, those with severe underlying diseases, or those enrolled in the study within the previous 12 months were not sampled.

Nasopharyngeal specimens were collected by trained paediatricians using nylon-tipped wire swabs. Swabs were inserted into the anterior nares, gently rubbed on the nasopharyngeal wall, removed, and immediately placed in transport medium (Copan Venturi Transystem®, Brescia, Italy). Samples were transferred within 48 hours to the National Reference Centre for Pneumococci (NRCP, https://cnr-pneumo.com/) or at Robert Debré Hospital (Paris, France) for analysis[1,3,4].

*Invasive pneumococcal disease*

In France, pneumococcal isolates collected from invasive pneumococcal disease (IPD), defined as the isolation of pneumococci from a normally sterile site, in children are sent to the NRCP for serotyping and antimicrobial susceptibility testing[7]. Previous capture-recapture studies estimated that this surveillance system for IPD by *Santé publique France*[8], involving more than 250 hospital laboratories, covered 73% to 79% of inpatient hospital stays[7]. Data collected included age, sex, participating centre, site and date of isolation, and clinical presentation.

*Microbiological investigations*

Each nasopharyngeal swab was swirled in 200 µL of brain-heart infusion, and 20 µL was smeared onto Columbia blood agar supplemented with colistin and nalidixic acid. Bacterial isolates were identified based on colony morphology and Gram staining. At least 5 colonies with the same morphology were serotyped together. If *S. pneumoniae* colonies displayed different morphologies, the different colonies were subcultured and then serotyped. In addition, the Statens Serum Institute guidelines were followed for serotyping. Pooled antisera were used according to a chessboard scheme that allowed for a detection of a mix of serogroups. Serotyping was performed by using latex particles sensitized with antisera purchased from the Statens Serum Institute (Copenhagen, Denmark)[9].

*S. pneumoniae* antimicrobial susceptibility profile was determined by minimal inhibitory concentration (MIC) using the agar-dilution method between 2001 and 2016 and with the broth microdilution method between 2017 and 2023. Non-susceptibility to penicillin G was defined as MIC >0.064 mg/L, and to amoxicillin, cefotaxime and erythromycin as MIC >0.5 mg/L, according to European Committee on Antimicrobial Susceptibility Testing (EUCAST) guidelines[10]. Pneumococcal isolates were preserved in cryotubes at -80°C at the NRCP.

**Supplementary Methods 2 | Pneumococcal conjugate vaccines and study periods.**

*National immunization program*

In 2001, 7-valent pneumococcal conjugate vaccine (PCV7) was licensed in France but not covered by French health insurance schemes, resulting in low vaccine coverage (<10% until December 2002). In January 2003, PCV7 was reimbursed and recommended for children at risk (i.e., children breastfed for less than 2 months, children in daycare with two or more other children, or children in families with more than two children). These criteria led to a 58% PCV7 coverage in 9-month-old infants in 2005. In June 2006, the recommendation and reimbursement were extended to all children younger than 2 years, leading to a 78% vaccine coverage in 2007. In 2010, PCV7 was replaced by PCV13 for all children younger than 2 years (two primary doses at ages 2 and 4 months and a booster at age 11 months), without catch up. Since 2011, PCV13 coverage for the primary vaccination series in infants aged 9 months has been greater than 91%[11]. PCV15 and PCV20 were not implemented in France during the study period.

*Study periods*

Based on the introduction timelines and vaccine coverage of PCV7 and PCV13 in France, we selected isolates collected during four distinct epidemiological years representative of successive vaccine eras: 1) the pre-PCV7 period in 2002–2003, with minimal vaccine coverage; 2) the PCV7 period in 2008–2009; 3) the early PCV13 period in 2012–2013, corresponding to the widespread adoption of PCV13; and 4) the late PCV13 period in 2022–2023.

For each period, all isolates collected during the winter season from nasopharyngeal carriage and IPD were included and subjected to whole-genome sequencing. Rather than sequencing a random subset across broad time windows, we selected all pneumococcal isolates collected from children in the national surveillance network for each study year. This exhaustive, year-by-year sampling ensured full representation of circulating strains within each period and enabled precise estimation of carriage rate and IPD incidence.

**Supplementary Methods 3 | Whole genome analysis.**

DNA was extracted from single colonies using Chemagic DNA/RNA Viral Kit on a Chemagic Prime instrument (Revvity, Waltham, MA, USA). Libraries were prepared using DNAPrep (Illumina, San Diego, Unites States) libraries on a Fluent 780 DreamPrep system (Tecan, Männedorf, Switzerland). Pair-end sequencing (2 × 150 bp) was performed on an Illumina NovaSeq™ 6000 system (Illumina®, San Diego, USA) using an SP v1.5 300-cycle kit.

Paired-end reads were adapter- and quality-trimmed using Trim Galore v0.6.7 (Cutadapt v3.7) with a Phred quality threshold of 20 and a minimum read length of 50 bp. Trimmomatic (v0.38) was used to remove residual low-quality bases from read termini (LEADING:20, TRAILING:20), performed sliding window (SLIDINGWINDOW:15:20), and exclusion of short reads (MINLEN:50).

Sequencing read quality was assessed using FastQC v0.11.8, and results were summarised using MultiQC v1.7. Theoretical sequencing depth was estimated after trimming from the total number of sequenced bases using a reference genome size of 2.1 Mb for *S. pneumoniae*[12]. Taxonomic profiling of trimmed paired-end reads was performed using MetaPhlAn v4.0.2 with the CHOCOPhlAn marker gene database (mpa_vJan21_CHOCOPhlAnSGB_202103).

Assembly was performed using SPAdes v3.15.4. Genome assembly statistics were calculated using QUAST v5.0.2. Genome size and contiguity metrics (N50 and L50) were calculated using contigs ≥500 bp, corresponding to the default reporting threshold in QUAST.

Serotypes were inferred using SeroBA v2.0.5[13]. In silico predicted serotypes were compared with in vitro data, and discordance findings led to repeat in vitro serotyping. Global Pneumococcal Sequence Clusters (GPSCs) were assigned with PopPUNK v2.7.5[14], using core and accessory distances, calculated by shared DNA k-mers (reference database available at https://www.pneumogen.net/gps/#/). PopPunk database v.10 for *Streptococcus pneumoniae* was used (https://www.bacpop.org/poppunk-databases/).

De novo genome annotation was performed using Prokka v1.14 with default parameters unless otherwise specified, using the bacterial genetic code (translation table 11). Annotated genomes were used to infer the pan-genome using Roary[15], clustering orthologous genes at 95% amino-acid sequence identity. Core genes were defined as those present in at least 95% of genomes. Core gene sequences were aligned using MAFFT and concatenated into a single core genome alignment.

Phylogenetic reconstruction was performed using maximum-likelihood inference implemented in IQ-TREE v1.6.9[16], based on the concatenated core genome alignment. The GTR nucleotide substitution model was applied, and branch support was assessed using 1000 ultrafast bootstrap replicates and 1000 SH-like approximate likelihood ratio tests (SH-aLRT). Phylogenetic inference was conducted without a designated reference genome or outgroup. The resulting tree was midpoint-rooted for visualisation and interpretation of population structure and visualized using the Interactive Tree Of Life (iTOL) web server[17]. GPSC assignments inferred independently using PopPUNK were subsequently mapped onto the phylogeny to assess concordance between population structure and core-genome phylogenetic relationships. Monophyly of GPSCs was assessed from the core-genome maximum-likelihood phylogeny in R using the `is.monophyletic()` function in the *ape* package[18]. GPSCs represented by a single isolate were considered trivially monophyletic. For GPSCs classified as non-monophyletic, the number and size of phylogenetically distinct clades were quantified by identifying maximal "pure" internal nodes whose descendant tips all belonged to the same GPSC. Descendant tip sets for internal nodes were obtained using the `Descendants()` function implemented in the *phangorn* package[19].

### Supplementary Methods 4 | Model selection based on AIC comparison.

**Supplementary Table S1 | Model comparison for serotype group carriage rates and IPD incidence.**

| Dataset/ model specification | Polynomial degree (time) | AIC | ΔAIC |
|---|---|---|---|
| **Carriage rate of** | | | |
| **PCV7 serotypes** | | | |
| Time independent | 0 | 1914.78 | 1776.98 |
| Linear with time | 1 | 374.43 | 236.62 |
| Quadratic with time* | 2 | 137.81 | 0 |
| Cubic with time | 3 | 139.80 | 1.99 |
| **PCV13 non-PCV7 serotypes** | | | |
| Time independent | 0 | 1069.04 | 844.55 |
| Linear with time | 1 | 458.04 | 233.55 |
| Quadratic with time | 2 | 327.29 | 102.80 |
| Cubic with time* | 3 | 224.49 | 0 |
| **Non-PCV13 serotypes** | | | |
| Time independent | 0 | 1648.99 | 1449.82 |
| Linear with time | 1 | 421.76 | 222.59 |
| Quadratic with time* | 2 | 199.17 | 0 |
| Cubic with time | 3 | 200.25 | 1.07 |
| **IPD incidence of** | | | |
| **PCV7 serotypes** | | | |
| Time independent | 0 | 3468.80 | 3242.45 |
| Linear with time | 1 | 362.72 | 136.37 |
| Quadratic with time | 2 | 284.01 | 57.66 |
| Cubic with time* | 3 | 226.34 | 0 |
| **PCV13 non-PCV7 serotypes** | | | |
| Time independent | 0 | 3356.66 | 2902.59 |
| Linear with time | 1 | 1751.21 | 1297.15 |
| Quadratic with time | 2 | 897.98 | 443.91 |
| Cubic with time* | 3 | 454.07 | 0 |
| **Non-PCV13 serotypes** | | | |
| Time independent | 0 | 905.97 | 622.56 |
| Linear with time | 1 | 408.44 | 125.04 |
| Quadratic with time | 2 | 308.75 | 25.34 |
| Cubic with time* | 3 | 283.41 | 0 |

* Selected polynomial degree based on AIC minimization. For each dataset, linear, quadratic and cubic polynomial specifications of calendar year were compared, and the model with the lowest AIC was retained, favouring the most parsimonious specification when ΔAIC < 2. ΔAIC was calculated as the difference between the AIC of each model and the minimum AIC across models ($\Delta AIC = AIC_i - AIC_{min}$).
PCV, pneumococcal conjugate vaccine; AIC, Akaike information criterion.

**Supplementary Table S2 | Model comparison for GPSC groups in serotype-fixed and lineage-fixed scenarios.**

| Dataset/ model specification | Polynomial degree (time) | AIC | ΔAIC |
|---|---|---|---|
| **Serotype-fixed scenario** | | | |
| **PCV7-type GPSCs** | | | |
| Time independent + mean log(MIC) | 0 | 736.13 | 533.83 |
| Linear with time + mean log(MIC) | 1 | 227.59 | 25.29 |
| Quadratic with time + mean log(MIC)* | 2 | 202.31 | 0 |
| Cubic with time + mean log(MIC) | 3 | 203.55 | 1.24 |
| **PCV13 non-PCV7-type GPSCs** | | | |
| Time independent + mean log(MIC) | 0 | 555.20 | 228.47 |
| Linear with time + mean log(MIC) | 1 | 404.92 | 78.18 |
| Quadratic with time + mean log(MIC) | 2 | 350.33 | 23.60 |
| Cubic with time + mean log(MIC)* | 3 | 326.74 | 0 |
| **Non-PCV13-type GPSCs** | | | |
| Time independent + mean log(MIC) | 0 | 925.31 | 219.00 |
| Linear with time + mean log(MIC) | 1 | 786.57 | 80.26 |
| Quadratic with time + mean log(MIC)* | 2 | 706.31 | 0 |
| Cubic with time + mean log(MIC) | 3 | 707.48 | 1.17 |
| **Lineage-fixed scenario** | | | |
| **PCV7-type GPSCs** | | | |
| Time independent + mean log(MIC)* | 0 | 830.37 | 0 |
| Linear with time + mean log(MIC) | 1 | 832.35 | 1.98 |
| Quadratic with time + mean log(MIC) | 2 | 834.33 | 3.96 |
| Cubic with time + mean log(MIC) | 3 | 836.30 | 5.93 |
| **PCV13 non-PCV7-type GPSCs** | | | |
| Time independent + mean log(MIC)* | 0 | 780.76 | 0 |
| Linear with time + mean log(MIC) | 1 | 782.41 | 1.65 |
| Quadratic with time + mean log(MIC) | 2 | 784.40 | 3.64 |
| Cubic with time + mean log(MIC) | 3 | 786.40 | 5.64 |
| **Non-PCV13-type GPSCs** | | | |
| Time independent + mean log(MIC)* | 0 | 876.25 | 0 |
| Linear with time + mean log(MIC) | 1 | 878.24 | 2.00 |
| Quadratic with time + mean log(MIC) | 2 | 878.89 | 2.65 |
| Cubic with time + mean log(MIC) | 3 | 880.86 | 4.62 |

*Selected polynomial degree based on AIC minimization. For each dataset, time-independent, linear, quadratic and cubic polynomial specifications of calendar year were compared, and the model with the lowest AIC was retained, favouring the most parsimonious specification when ΔAIC < 2. ΔAIC was calculated as the difference between the AIC of each model and the minimum AIC across models ($\Delta AIC = AIC_i - AIC_{min}$).
Mean log(MIC) corresponds to the mean log-transformed MIC of isolates within each GPSC in each period.
GPSCs were grouped according to their pre-PCV7 (2002–2003) serotype composition.
AIC, Akaike information criterion; GPSC, Global Pneumococcal Sequence Cluster; PCV, pneumococcal conjugate vaccine; MIC, minimum inhibitory concentration.

**Supplementary Methods 5 | Construction of counterfactual scenarios.**

*Numerical optimisation of GPSC carriage rates under the serotype-fixed scenario*

Let us define $f_g(S,t)$ the observed relative proportion of serotype group $S$ (PCV7, PCV13 non-PCV7, non-PCV13) in GPSC $g$ in year *t*, $C(g,t)$ the total carriage rate of GPSC *g* in year *t* and $C(S,t)$ the total carriage rate of serotype group *S* in year *t*, such that:

$$C(S,t) = \sum_g C(g,t) f_g(S,t)$$

Under the serotype-fixed scenario, we fixed the counterfactual relative proportion of serogroup $S$ in GPSC $g$ $f'_g(S) = f_g(S, 2002)$ constant over time, at its observed pre-PCV7 level. Hence, the counterfactual total carriage rate of serotype group *S* in year *t,* computed in the same way as the observed one:

$$C'(S,t) = \sum_g C'(g,t) f'_g(S)$$

with $C'(g,t)$ such that the counterfactual carriage rate of GPSC *g* in year *t*. We estimated GPSC carriage rates $C'(g,t)$ at year *t* through numerical optimization based on the Nelder-Mead algorithm[20] by using the *optim* function in R. The function minimized was the sum of squares of the differences between $C(S,t)$ and $C'(S,t)$ for the three serotype groups:

$$\sum_S (C(S,t) - C'(S,t))^2$$

The initial values of $C'(g,t)$ were the observed carriage rates $C(g,t)$ in the same year. We performed the optimization under two additional constraints: that $C'(g,t) \geq 0$ for any *g* and *t* and that the sum of all counterfactual carriage rates was equal to the observed total carriage rate, i.e that:

$$\sum_g C'(g,t) = \sum_g C(g,t)$$

To appropriately perform binomial regressions on the carriage rates, the counterfactual carriage rates $C^{ser}(g,t)$ actually considered were adjusted so that the product of the rates by the number $N_t$ of samples in year *t* was integer:

$$C^{ser}(g,t) = \frac{round\left(N_t\, C'(g,t)\right)}{N_t}$$

**Supplementary Methods 6 | Quantification of mechanism contribution.**

Let us define $Y_G^{ser}(t)$, $Y_G^{lin}(t)$ and $Y_G^{obs}(t)$ the carriage rates of GPSC group *G* in year *t* estimated respectively by the regression on the counterfactual data of the serotype-fixed scenario, the regression on the counterfactual data of the lineage-fixed scenario, and the regression on the observed data. The weighted combination of the estimates from the two scenarios can be expressed as:

$$Y_G'(t) = w_G Y_G^{lin}(t) + (1 - w_G) Y_G^{ser}(t)$$

with $w_G \in [0,1]$ a factor describing the relative contribution of the lineage-fixed scenario, $w_G = 0$, $w_G = 1$ and $w_G = 0.5$ respectively representing contribution of the lineage-fixed scenario only, contribution of the serotype-fixed scenario only and an equal contribution of both scenarios.

We estimated $w_G$ through numerical optimization based on the Nelder-Mead algorithm[20] by using the *optim* function in R. The function minimized was the sum of squared differences between the weighted combination of estimates from the two scenarios and the estimate from the data-informed model across all years (2002–2023):

$$\sum_t \left(Y_G'(t) - Y_G^{obs}(t)\right)^2$$

We estimated separately four values of $w_G$, one per GPSC group (PCV7-type, PCV13 non-PCV7-type, non-PCV13-type) and one for all GPSCs. For the latter, the sum of the estimates for each GPSC group was used, weighted by their respective relative frequency in year *t*, such that the function minimized was:

$$\sum_t \left( \frac{\sum_{G \in \{\text{PCV7,PCV13,} \atop \text{non-PCV13}\}} \left( Y_{\text{G}}^{obs}(t) \left( Y_{\text{G}}'(t) - Y_{\text{G}}^{obs}(t) \right)^2 \right)}{Y_{\text{PCV7}}^{obs}(t) + Y_{\text{PCV13}}^{obs}(t) + Y_{\text{non-PCV13}}^{obs}(t)} \right)$$

The regressions on observed and counterfactual data each provided a model estimate and a standard error for the carriage rate of each GPSC group in each year. We used parametric simulation based on these estimates to generate 1,000,000 sets of estimates for each model and obtained a distribution of values of $w_G$ for each PCV group and for all GPSCs. We derived 95% confidence intervals from these distributions, their 2.5$^{th}$ and 97.5$^{th}$ percentiles corresponding to the boundaries of the interval.

**Supplementary Methods 7 | Estimation of serotype-specific invasive pneumococcal disease from carriage scenarios.**

We computed the incidence rate of IPD of serotype *s* based on its carriage rate in year *t*, noted $C(s,t)$, as follows:

$$\hat{I}(s,t) = k\, C(s,t)\, DP_s$$

with $DP_s$ the serotype-specific invasive disease potential and $k$ a calibration constant, which was defined such that the sum of $\hat{I}(s,t)$ for all serotypes matched the total IPD incidence observed in 2002, noted $I(2002)$:

$$k = \frac{I(2002)}{\sum_s C(s,2002)\, DP_s} = 501.32$$

We assigned each serotype represented in our carriage dataset a single, time-invariant value of $DP_s$, based on four sources of carriage–disease comparisons in paediatric populations[21–24]. When multiple estimates were available for a given serotype, we selected a single value a priori to ensure a unique value per serotype across the study period. We assigned a $DP_s$ of 1 to serotypes with no available published estimate and a $DP_s$ of 0.2 to non-typeable isolates, based on published estimates. The full list of DP values used in the analyses is provided in **Supplementary Table S3**.

**Supplementary Table S3 | Full list of serotype-specific invasive disease potential estimates.**

| Serotype | DP |
|---|---|
| 1 | 33.4 |
| 8 | 10.9 |
| 5 | 9.0 |
| 12F | 7 |
| 7F | 7 |
| 24F | 6.6 |
| 3 | 5.5 |
| 18A | 5.4 |
| 33F | 5 |
| 25A/38 | 4.5 |
| 4 | 3.7 |
| 14 | 3.4 |
| 6B | 3.1 |
| 22F | 3 |
| 31 | 3 |
| 19A | 2.4 |
| 9V | 2.4 |
| 18C | 2.1 |
| 11B* | 1 |
| 15F* | 1 |
| 16A* | 1 |
| 18B* | 1 |
| 18F* | 1 |
| 19B* | 1 |
| 20* | 1 |
| 27* | 1 |
| 28A* | 1 |
| 29* | 1 |
| 35F* | 1 |
| 37* | 1 |
| 7B* | 1 |
| 9L* | 1 |
| 17F | 0.7 |
| 9N | 0.6 |
| 16F | 0.5 |
| 19F | 0.5 |
| 23F | 0.8 |
| 35B | 0.8 |
| 15A | 0.4 |
| 23B | 0.4 |
| 10A | 0.3 |
| 15B/C | 0.3 |
| 6A/C | 0.3 |
| NT | 0.2 |
| 10B | 0.2 |
| 34 | 0.1 |
| 11A | 0 |
| 13 | 0 |
| 21 | 0 |
| 23A | 0 |

*Serotypes with no available published DP estimate.
DP, disease potential; NT, non-typable.

We first evaluated its validity on the observed serotype carriage data across the entire study period (2002–2023). The predicted IPD incidence closely reproduced the temporal trends observed in surveillance data (**Supplementary Fig. S1**).

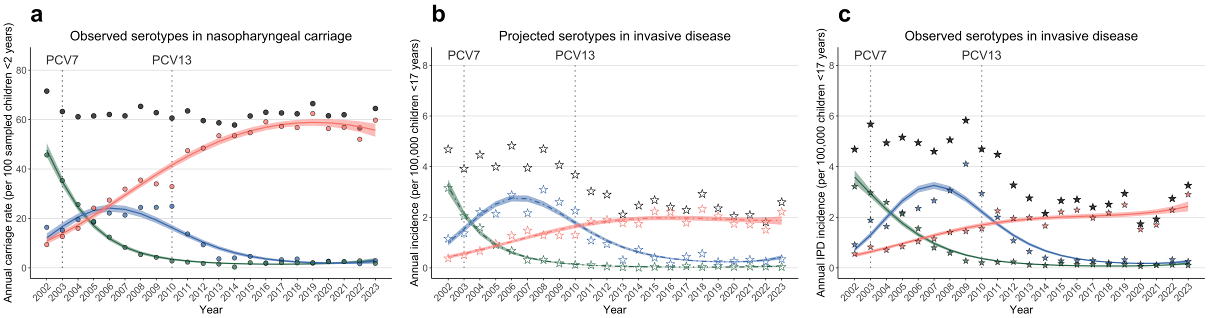


**Supplementary Fig. S1 | Predicted pneumococcal invasive disease from serotype carriage data and comparison with observed data. a–c,** Annual carriage rates in children aged ≤2 years (**a**), predicted IPD incidence (**b**), and observed IPD incidence (**c**) from 2002 to 2023 for PCV7 serotypes (green), PCV13 non-PCV7 serotypes (blue), and non-PCV13 serotypes (red). Dots and stars represent observed data (dots for carriage, stars for IPD), and curves represent model fits with 95% CI (shaded bands). Open stars indicate predicted IPD incidence, whereas filled stars indicate observed IPD incidence. Rates are expressed per 100 children for carriage and per 100,000 children aged ≤17 years for IPD. IPD incidence of serotypes was estimated as proportional to the product of carriage and serotype-specific invasive disease potential derived from the literature[21–24]. Vertical dashed lines indicate the progressive implementation of PCV7 from 2003 and the introduction of PCV13 in 2010.
PCV, pneumococcal conjugate vaccine; IPD, invasive pneumococcal disease.

We then applied the same method to the counterfactual carriage estimates derived from the two mechanistic scenarios.

Under the serotype-fixed scenario, we computed the carriage rate of serotype *s* in year *t* as follows:

$$C^{ser}(s,t) = \sum_{g} C^{ser}(g,t)\, f_g(s, 2002)$$

with $C^{ser}(g,t)$ the carriage rate of GPSC *g* in year *t* and $f_g(s, 2002)$ the pre-PCV7 proportion of serotype *s* within GPSC *g*.

Under the lineage-fixed scenario, we computed the carriage rate of serotype *s* in year *t* as follows:

$$C^{lin}(s,t) = \sum_{g} C(g, 2002) \frac{Q_g(s,t)}{\sum_s Q_g(s,t)}$$

with $C(g, 2002)$ the pre-PCV7 carriage rate of GPSG *g* and $Q_g(s,t)$ the non-scaled counterfactual proportion of serotype *s* in GPSC *g*, which was computed as follows:

$$Q(s,t) = f_g(s, 2002) \frac{C(s,t)}{C(s, 2002)}$$

with $f_g(s, 2002)$, the pre-PCV7 proportion of serotype *s* within GPSC *g* and $C(s,t)$ the total carriage rate of serotype *s* in year *t*. Hence, $Q(s,t)$ was to the product between the within-lineage proportion serotype of *s* at baseline and the ratio of overall change in serotype *s* compared to baseline. The resulting values was normalized to maintain an overall lineage carriage rate identical to its observed carriage at baseline.

We assessed scenario performance with Pearson's correlation coefficients comparing Log of serotype-specific IPD incidence based on counterfactual data with the Log of IPD incidence rates from surveillance data, both in the most recent study period (late PCV13 period, 2022–2023). To include serotypes whose incidence estimates were zero in counterfactual or observed data, we assigned them a value of $0.5/N_t'$, with $N_t'$ the total number of IPD observations in the corresponding period.

### Supplementary Results.

#### Supplementary Results 1 | Whole genome quality control.

*Read quality assessment*

MultiQC summaries indicated consistently high sequencing quality across samples, with all samples passing per-base sequence quality thresholds and efficient removal of adapter sequences. Warnings related to per-sequence GC content and sequence duplication levels were frequent and are consistent with the narrow GC distribution and high sequencing depth expected for bacterial whole-genome sequencing. No systematic quality failures indicative of contamination or technical artefacts were observed (**Supplementary Table S4**).

**Supplementary Table S4 | Summary of FastQC quality control results across samples.**

| FastQC module | Pass, n (%) | Warn, n (%) | Fail, n (%) |
|---|---|---|---|
| Adapter content | 4802 (100) | 0 | 0 |
| Over-represented sequences | 4800 (9.99) | 2 (0.04) | 0 |
| Per-base sequence quality | 4802 (100) | 0 | 0 |
| Per-sequence GC content | 573 (11.9) | 4229 (88.1) | 0 |
| Sequence duplication levels | 479 (10.0) | 1817 (37.8) | 2506 (52.2) |

For each FastQC module, the table reports the number of samples (n) and corresponding percentages (%) classified as pass, warn, or fail. Results were aggregated across all samples using MultiQC v1.7. Samples were flagged as fail for sequence duplication levels when fewer than 20% of reads were unique, and for GC content when the deviation from the theoretical distribution exceeded 30%.

Based on read counts and average read length, the median theoretical sequencing depth after trimming was 352x (IQR, 231–471x), with all samples exceeding 35x coverage. This depth is consistent with high-quality bacterial genome assembly and explains the elevated sequence duplication levels observed in read-level quality control.

*Relationship between read-level QC and assembly quality*

Despite FastQC warnings related to GC content and sequence duplication levels, assembly quality metrics assessed using QUAST were consistent across samples, with stable genome sizes, low fragmentation and high N50 values, indicating that these read-level warnings did not adversely affect genome assembly quality (**Supplementary Fig. S2**).

*Genome assembly quality*

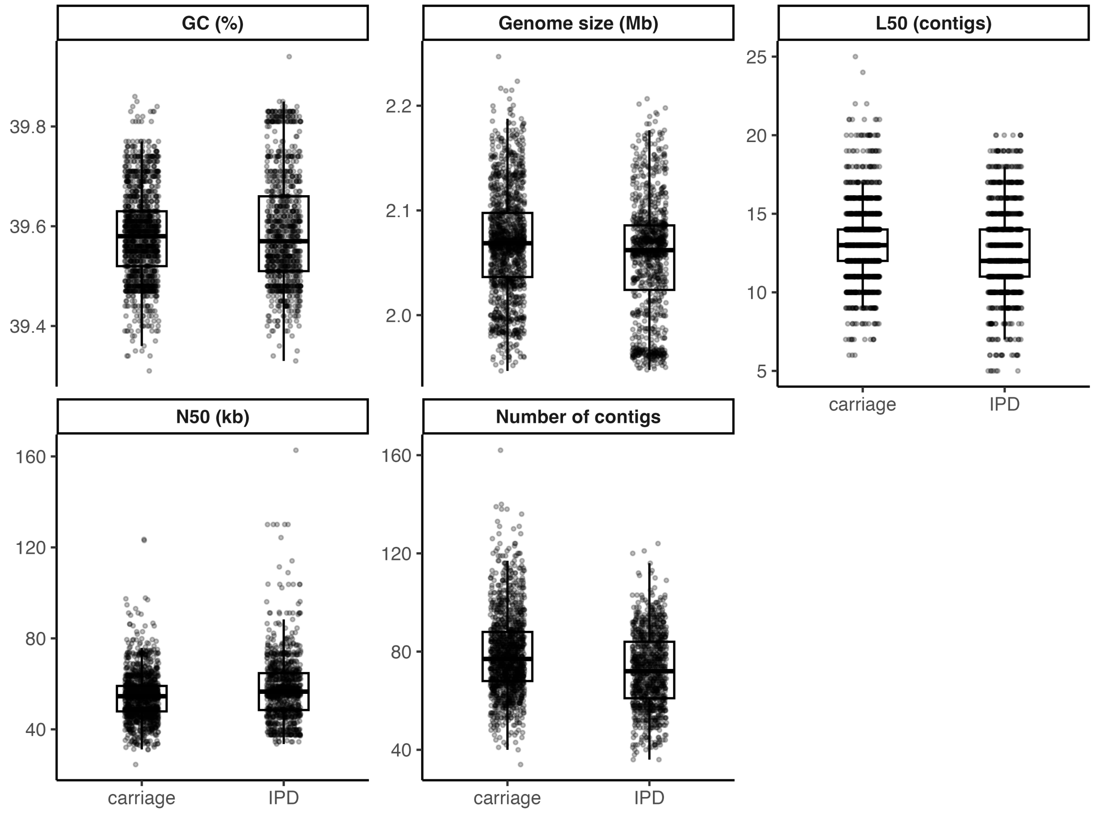


**Supplementary Data Fig. S2 | Genome assembly quality metrics. Boxplots show the distribution of genome size, GC content, number of contigs, N50 and L50 for carriage and IPD isolates**. Points represent individual genome assemblies. Genome size, N50 and L50 were calculated using contigs retained by QUAST for assembly quality assessment. Assemblies comprised 34-162 contigs (median, 75). N50 values ranged from 24,526 to 162,714 bp (median [IQR], 55,077 [48,286–61,758]), with corresponding L50 values of 5–25 contigs (median [IQR], 13 [11–14]). GC content ranged from 39.3% to 39.9%. All assembly statistics were computed by QUAST using contigs ≥500 bp.

*Taxonomic validation*

MetaPhlAn profiling confirmed the taxonomic purity of the sequencing data, with a median of 100% of reads assigned to *S. pneumoniae* (minimum 99.18%). No other species exceeded 1% relative abundance in any sample (maximum observed, 0.82%).

**Supplementary Results 2 | Pneumococcal population structure defined by GPSCs, serotypes and core-genome phylogeny.**

*GPSC distribution*

Among the 95 GPSC identified, 23 high-frequency GPSCs, each representing >1% of isolates, accounted for 1,983 isolates (82.6% of the dataset), whereas the remaining 72 low-frequency GPSCs accounted for 418 isolates (17.4%). The number of isolates assigned to each GPSC is shown in **Supplementary Fig. S3**.

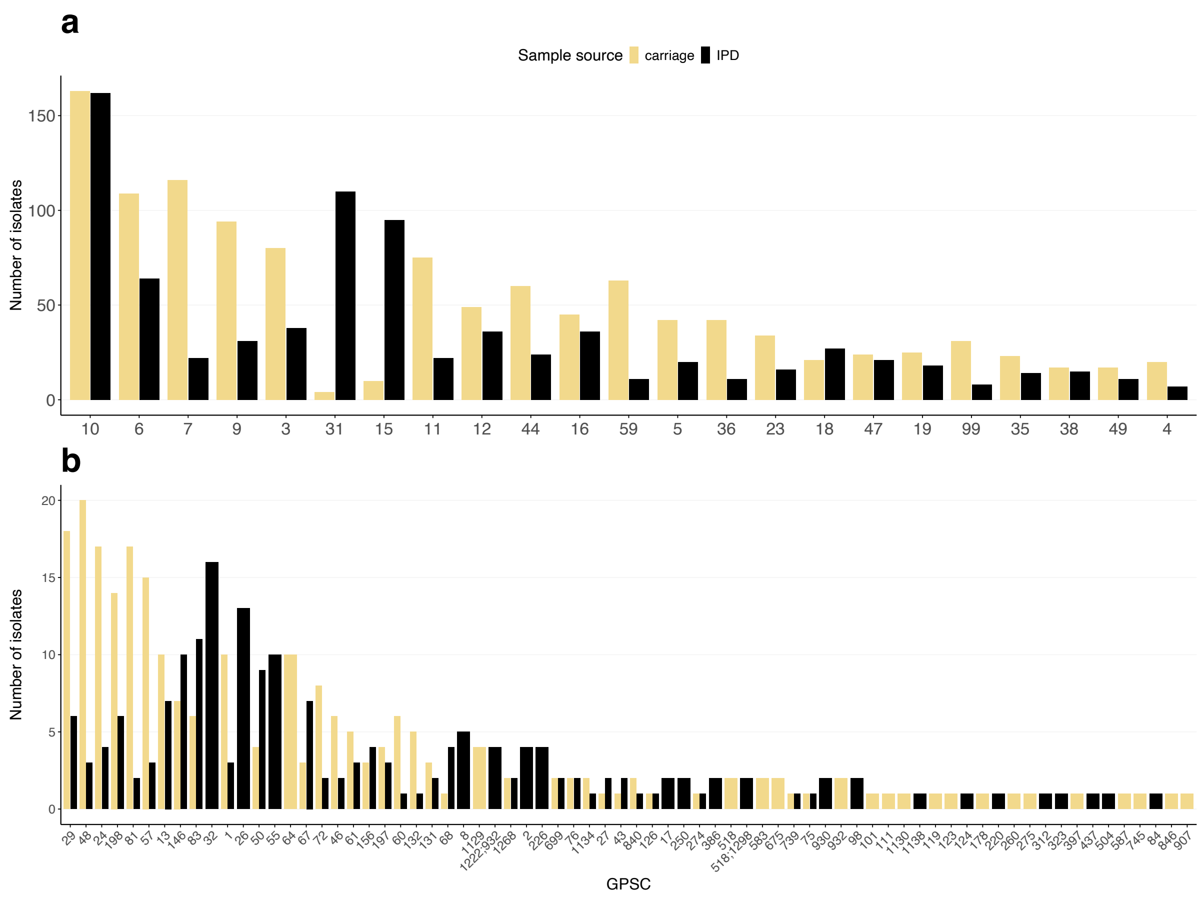


**Supplementary Fig. S3 | Distribution of GPSCs by isolate source**. Distribution of the 23 high-frequency GPSCs, each representing >1% of isolates (a), and 72 low-frequency GPSCs (each representing ≤1% of isolates) (b) among 2,401 *S. pneumoniae* genomes included in the genomic analysis. Bars indicate the number of isolates per GPSC, stratified by isolate source (carriage in yellow and IPD in black). GPSC, Global Pneumococcal Sequence Cluster; IPD, invasive pneumococcal disease.

*Serotype composition of GPSCs*

Serotype composition within each high-frequency GPSC is shown in **Supplementary Fig. S4**. The following GPSCs were associated with a single serotype: GPSC31-1, GPSC12-3, GPSC15-7F, GPSC35-10A, GPSC99-21, and GPSC19-22F. In contrast, most GPSCs encompassed multiple serotypes and exhibited heterogenous distributions between carriage and IPD isolates.

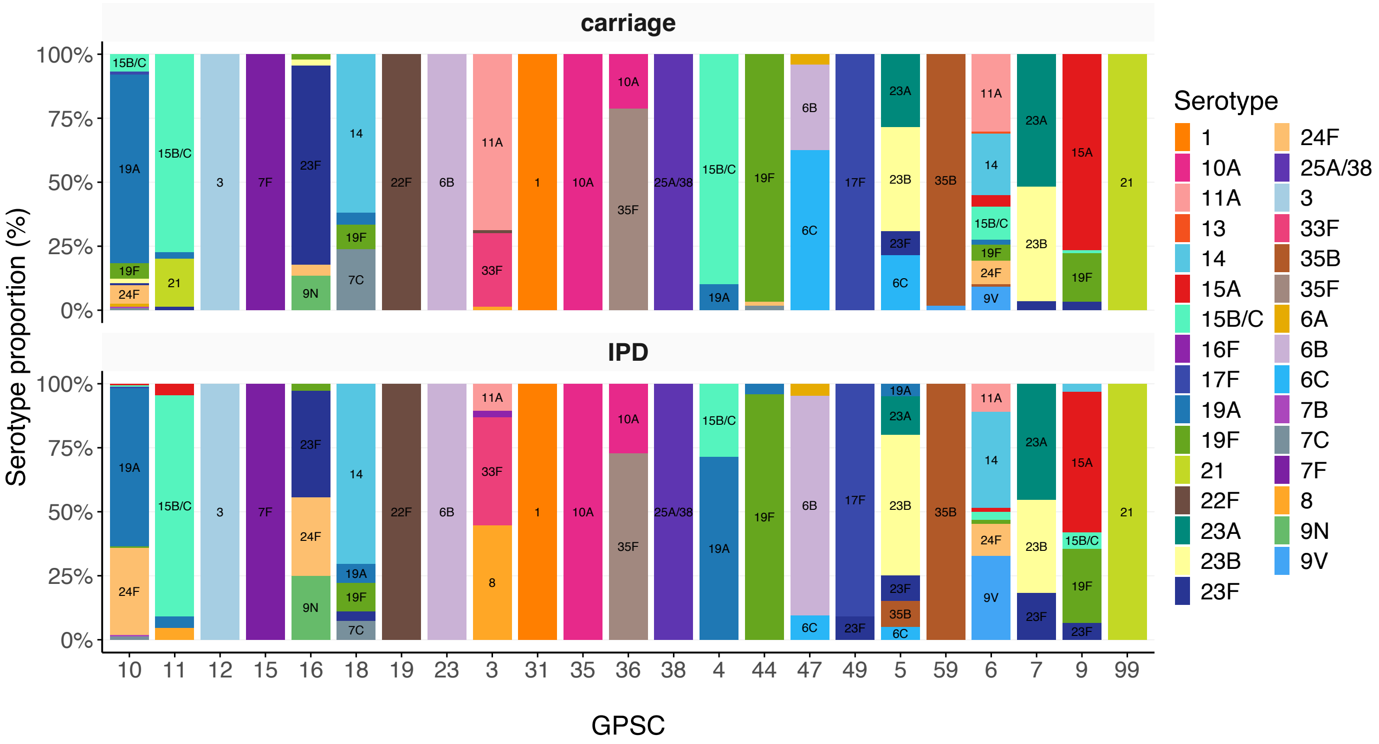


**Supplementary Fig. S4 | Serotype composition of high-frequency GPSCs by isolate source.** Stacked bar plots show the relative contribution of individual serotypes within high-frequency GPSCs (>1% of isolates), stratified by isolate source (carriage and IPD). Bars represent serotype proportions within each GPSC, and serotype labels are shown for components representing ≥5% of isolates within a GPSC.

*<u>Core-genome phylogeny and concordance with GPSC assignment</u>*

Core-genome phylogenetic analysis showed that 91 of 95 GPSCs formed monophyletic clades. Four GPSCs (GPSC10, GPSC518, GPSC518/1298, and GPSC1222/932), showed apparent non-monophyly, each because of one single isolate. For three of them, ambiguous composite PopPUNK assignments (GPSC518;1298 and GPSC1222;932) may have accounted for the observed non-monophyletic patterns, rather than structured phylogenetic discordance.

GPSC10 was characterised by a single large clade comprising 324 of 325 isolates, with one phylogenetically incongruent carriage isolate assigned to GPSC907 and embedded within the GPSC10 clade. The isolate was classified as non-encapsulated (non-typeable by in vitro serotyping) and carried an NCC1-type non-capsular locus encoding the surface protein PspK, as identified by seroBA. Non-encapsulated pneumococci are predominantly associated with nasopharyngeal carriage rather than IPD, and loss of capsule expression with replacement of the capsular locus by a non-capsular locus encoding PspK has been shown to enhance epithelial adherence and persistence during colonisation.[25,26] The phylogenetic embedding of this isolate within an otherwise encapsulated GPSC10 clade is therefore consistent with a lineage-specific capsular loss event rather than misclassification, highlighting the genomic plasticity of pneumococci during carriage. Notably, the isolate was positioned at the interface of several GPSC10 subclades, spanning multiple internal clade boundaries, a pattern compatible with the high recombination rates observed during carriage, particularly among non-encapsulated pneumococci (**Supplementary Fig. S5**).

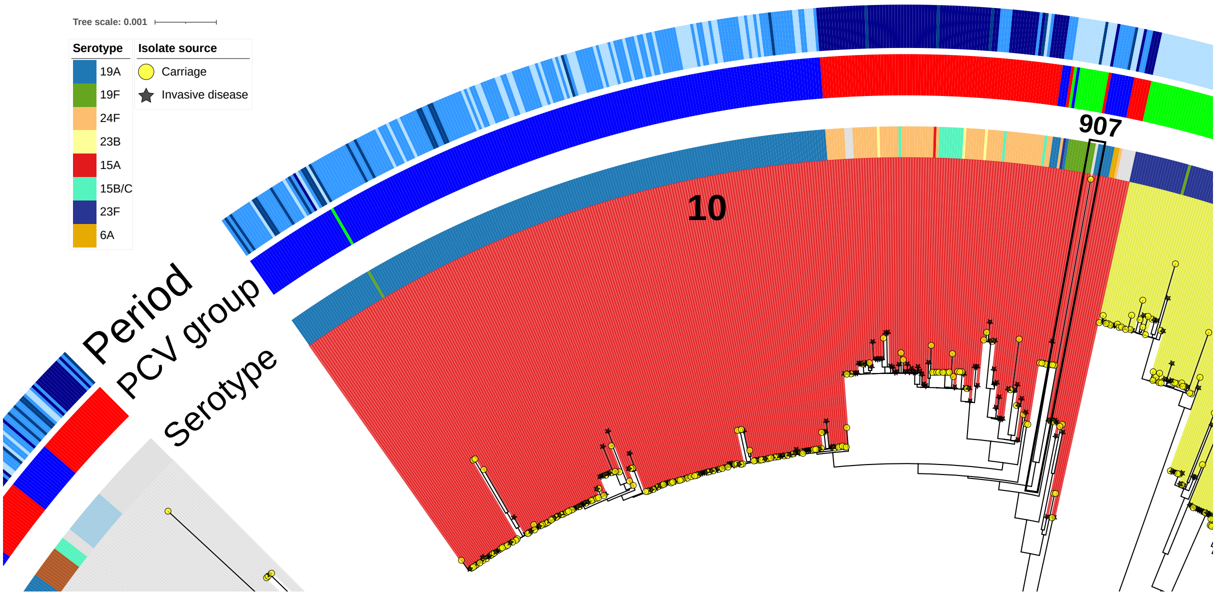


**Supplementary Fig. S5 | Core-genome phylogeny highlighting GPSC10. Midpoint-rooted maximum-likelihood phylogeny inferred from a concatenated core-genome alignment of 2401 *S. pneumoniae* isolates.** Tips represent individual isolates and are annotated by isolate source (carriage, yellow circles; invasive disease, black stars). Concentric colour strips from inner to outer indicate serotype, serotype PCV-group, and PCV-related sampling period. GPSC10 is shown as a dominant monophyletic clade (red), comprising 324 of 325 isolates. A single phylogenetically incongruent, non-encapsulated carriage isolate assigned to GPSC907 is embedded within the GPSC10 clade, accounting for the apparent polyphyly of GPSC10.

### Supplementary Results 3 | Models for serotype carriage rate and IPD incidence over time.

**Supplementary Data Table S5 | Model coefficients for carriage rate and IPD incidence of serotype groups from 2002 to 2023.**

| Dataset | Coefficient* | Estimate | Standard error | *P* value |
|---|---|---|---|---|
| **PCV7 serotypes** | | | | |
| **Carriage rate£** | Intercept | -3.840 | 0.073 | <0.0001 |
| | Year | -1.042 | 0.050 | <0.0001 |
| | $Year^2$ | 0.789 | 0.047 | <0.0001 |
| **IPD incidence$** | Intercept | -13.244 | 0.064 | <0.0001 |
| | Year | -1.739 | 0.082 | <0.0001 |
| | $Year^2$ | 0.574 | 0.043 | <0.0001 |
| | $Year^3$ | 0.309 | 0.040 | <0.0001 |
| **PCV13 non-PCV7 serotypes** | | | | |
| **Carriage rate£** | Intercept | -2.297 | 0.053 | <0.0001 |
| | Year | -1.863 | 0.090 | <0.0001 |
| | $Year^2$ | -0.189 | 0.048 | <0.0001 |
| | $Year^3$ | 0.520 | 0.049 | <0.0001 |
| **IPD incidence$** | Intercept | -11.417 | 0.030 | <0.0001 |
| | Year | -2.061 | 0.053 | <0.0001 |
| | $Year^2$ | -0.355 | 0.029 | <0.0001 |
| | $Year^3$ | 0.668 | 0.031 | <0.0001 |
| **Non-PCV13 serotypes** | | | | |
| **Carriage rate£** | Intercept | -0.009 | 0.028 | 0.757 |
| | Year | 0.722 | 0.022 | <0.0001 |
| | $Year^2$ | -0.356 | 0.024 | <0.0001 |
| **IPD incidence$** | Intercept | -10.871 | 0.021 | <0.0001 |
| | Year | 0.203 | 0.037 | <0.0001 |
| | $Year^2$ | -0.211 | 0.020 | <0.0001 |
| | $Year^3$ | 0.110 | 0.021 | <0.0001 |

£ Carriage rate is per 100 sampled children aged ≤ 2 years per year.
$ IPD incidence is per 100,000 children aged ≤17 years living in France per year.
* Model coefficients are reported on the logit scale.
PCV, pneumococcal conjugate vaccine; IPD, invasive pneumococcal disease; CI, confidence interval.

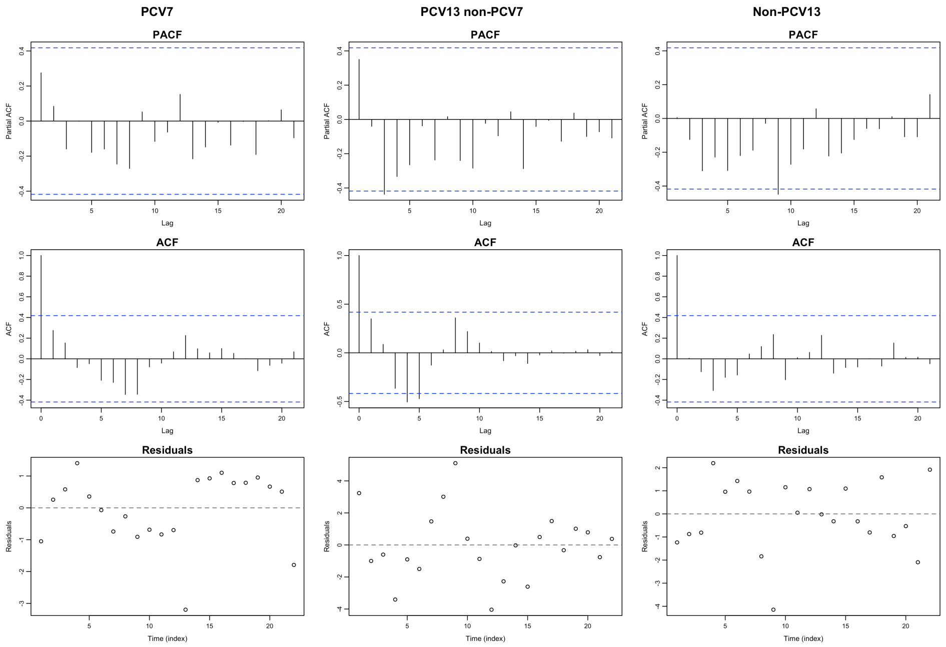


**Supplementary Fig. S6 | Goodness-of-fit diagnostics for the selected serotype carriage models.** Residual diagnostics for polynomial logistic regression models of annual serotype-group carriage rates in children aged ≤2 years. Columns correspond to PCV7, PCV13 non-PCV7 and non-PCV13 serotype groups. Rows show partial autocorrelation functions (PACF), autocorrelation functions (ACF) and residual time series. Blue dashed lines indicate approximate 95% confidence bounds. No substantial temporal autocorrelation or systematic residual structure was observed.

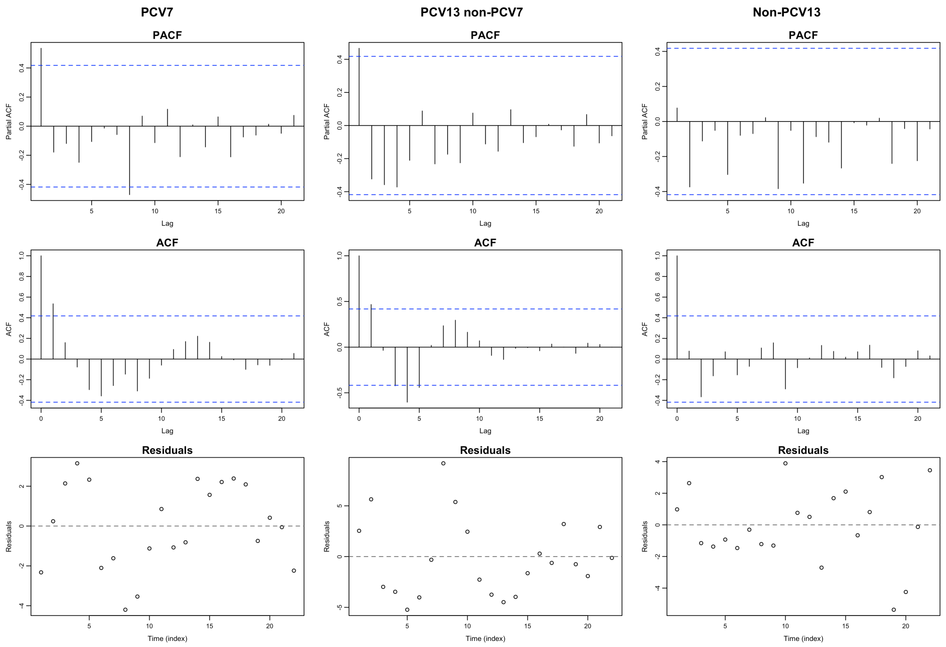


**Supplementary Fig. S7 | Goodness-of-fit diagnostics for the selected serotype IPD models.** Residual diagnostics for polynomial logistic regression models of annual serotype-group IPD incidence in children aged ≤17 years. Columns correspond to PCV7, PCV13 non-PCV7 and non-PCV13 serotype groups. Rows show partial autocorrelation functions (PACF), autocorrelation functions (ACF) and residual time series. Blue dashed lines indicate approximate 95% confidence bounds. No substantial temporal autocorrelation or systematic residual structure was observed.

### Supplementary Results 4 | GPSC characteristics and evolution in carriage.

**Supplementary Table S6 | Classification of GPSCs according to their serotype composition in the pre-PCV7 period (2002–2003).**

| | Number of isolates (%)* | | |
|---|---|---|---|
| **GPSC group/ GPSC** | **PCV7 serotypes** | **PCV13 non-PCV7 serotypes** | **Non-PCV13 serotypes** |
| **PCV7-type GPSCs** | | | |
| 1 | 7/7 (100.0) | 0/7 (0.0) | 0/7 (0.0) |
| 6 | 40/40 (100.0) | 0/40 (0.0) | 0/40 (0.0) |
| 7 | 4/5 (80.0) | 0/5 (0.0) | 1/5 (20.0) |
| 9 | 19/21 (90.5) | 0/21 (0.0) | 2/21 (9.5) |
| 16 | 36/37 (97.3) | 0/37 (0.0) | 1/37 (2.7) |
| 18 | 15/15 (100.0) | 0/15 (0.0) | 0/15 (0.0) |
| 23 | 32/32 (100.0) | 0/32 (0.0) | 0/32 (0.0) |
| 24 | 11/12 (91.7) | 1/12 (8.3) | 0/12 (0.0) |
| 43 | 1/1 (100.0) | 0/1 (0.0) | 0/1 (0.0) |
| 44 | 22/22 (100.0) | 0/22 (0.0) | 0/22 (0.0) |
| 47 | 8/10 (80.0) | 1/10 (10.0) | 1/10 (10.0) |
| 50 | 3/3 (100.0) | 0/3 (0.0) | 0/3 (0.0) |
| 68 | 1/1 (100.0) | 0/1 (0.0) | 0/1 (0.0) |
| 198 | 13/13 (100.0) | 0/13 (0.0) | 0/13 (0.0) |
| **PCV13 non-PCV7-type GPSCs** | | | |
| 10 | 11/46 (23.9) | 35/46 (76.1) | 0/46 (0.0) |
| 12 | 0/22 (0.0) | 22/22 (100.0) | 0/22 (0.0) |
| 13 | 0/4 (0.0) | 3/4 (75.0) | 1/4 (25.0) |
| 15 | 0/1 (0.0) | 1/1 (100.0) | 0/1 (0.0) |
| 29 | 0/9 (0.0) | 8/9 (88.9) | 1/9 (11.1) |
| 31 | 0/2 (0.0) | 2/2 (100.0) | 0/2 (0.0) |
| 64 | 0/5 (0.0) | 5/5 (100.0) | 0/5 (0.0) |
| 76 | 0/2 (0.0) | 2/2 (100.0) | 0/2 (0.0) |
| 197 | 0/3 (0.0) | 3/3 (100.0) | 0/3 (0.0) |
| 675 | 0/2 (0.0) | 2/2 (100.0) | 0/2 (0.0) |
| 739 | 0/1 (0.0) | 1/1 (100.0) | 0/1 (0.0) |
| 840 | 0/2 (0.0) | 2/2 (100.0) | 0/2 (0.0) |
| **Non-PCV13-type GPSCs** | | | |
| 3 | 0/17 (0.0) | 0/17 (0.0) | 17/17 (100.0) |
| 4 | 0/1 (0.0) | 0/1 (0.0) | 1/1 (100.0) |
| 5 | 4/9 (44.4) | 0/9 (0.0) | 5/9 (55.6) |
| 11 | 1/6 (16.7) | 2/6 (33.3) | 3/6 (50.0) |
| 19 | 0/2 (0.0) | 0/2 (0.0) | 2/2 (100.0) |
| 27 | 0/0 (0) | 0/0 (0) | 0/0 (0) |
| 35 | 0/1 (0.0) | 0/1 (0.0) | 1/1 (100.0) |
| 36 | 0/2 (0.0) | 0/2 (0.0) | 2/2 (100.0) |
| 38 | 0/5 (0.0) | 0/5 (0.0) | 5/5 (100.0) |
| 46 | 0/1 (0.0) | 0/1 (0.0) | 1/1 (100.0) |
| 48 | 0/8 (0.0) | 0/8 (0.0) | 8/8 (100.0) |
| 49 | 0/0 (0) | 0/0 (0) | 0/0 (0) |
| 57 | 0/1 (0.0) | 0/1 (0.0) | 1/1 (100.0) |
| 59 | 1/3 (33.3) | 0/3 (0.0) | 2/3 (66.7) |
| 60 | 0/0 (0) | 0/0 (0) | 0/0 (0) |
| 61 | 0/1 (0.0) | 0/1 (0.0) | 1/1 (100.0) |
| 67 | 1/2 (50.0) | 0/2 (0.0) | 1/2 (50.0) |
| 72 | 0/1 (0.0) | 0/1 (0.0) | 1/1 (100.0) |
| 75 | 0/0 (0) | 0/0 (0) | 0/0 (0) |
| 81 | 0/2 (0.0) | 0/2 (0.0) | 2/2 (100.0) |
| 83 | 0/0 (0) | 0/0 (0) | 0/0 (0) |
| 99 | 0/1 (0.0) | 0/1 (0.0) | 1/1 (100.0) |
| 101 | 0/0 (0) | 0/0 (0) | 0/0 (0) |
| 111 | 0/1 (0.0) | 0/1 (0.0) | 1/1 (100.0) |
| 119 | 0/0 (0) | 0/0 (0) | 0/0 (0) |
| 123 | 0/0 (0) | 0/0 (0) | 0/0 (0) |
| 126 | 0/1 (0.0) | 0/1 (0.0) | 1/1 (100.0) |
| 131 | 0/0 (0) | 0/0 (0) | 0/0 (0) |
| 132 | 0/0 (0) | 0/0 (0) | 0/0 (0) |
| 146 | 0/0 (0) | 0/0 (0) | 0/0 (0) |
| 156 | 0/0 (0) | 0/0 (0) | 0/0 (0) |
| 178 | 0/0 (0) | 0/0 (0) | 0/0 (0) |
| 260 | 0/0 (0) | 0/0 (0) | 0/0 (0) |
| 274 | 0/0 (0) | 0/0 (0) | 0/0 (0) |
| 275 | 0/0 (0) | 0/0 (0) | 0/0 (0) |
| 397 | 0/0 (0) | 0/0 (0) | 0/0 (0) |
| 518 | 0/0 (0) | 0/0 (0) | 0/0 (0) |

| | | | |
|---|---|---|---|
| 583 | 0/0 (0) | 0/0 (0) | 0/0 (0) |
| 587 | 0/0 (0) | 0/0 (0) | 0/0 (0) |
| 699 | 0/1 (0.0) | 0/1 (0.0) | 1/1 (100.0) |
| 745 | 0/0 (0) | 0/0 (0) | 0/0 (0) |
| 846 | 0/0 (0) | 0/0 (0) | 0/0 (0) |
| 907 | 0/0 (0) | 0/0 (0) | 0/0 (0) |
| 932 | 0/0 (0) | 0/0 (0) | 0/0 (0) |
| 1129 | 0/3 (0.0) | 0/3 (0.0) | 3/3 (100.0) |
| 1130 | 0/0 (0) | 0/0 (0) | 0/0 (0) |
| 1134 | 1/2 (50.0) | 0/2 (0.0) | 1/2 (50.0) |
| 1268 | 0/2 (0.0) | 0/2 (0.0) | 2/2 (100.0) |

*Number (%) of isolates belonging to each serotype group within each GPSC in the pre-PCV7 period (2002–2003).
GPSC, Global Pneumococcal Sequence Cluster; PCV, pneumococcal conjugate vaccine.

*Penicillin non-susceptibility of GPSCs across study periods*

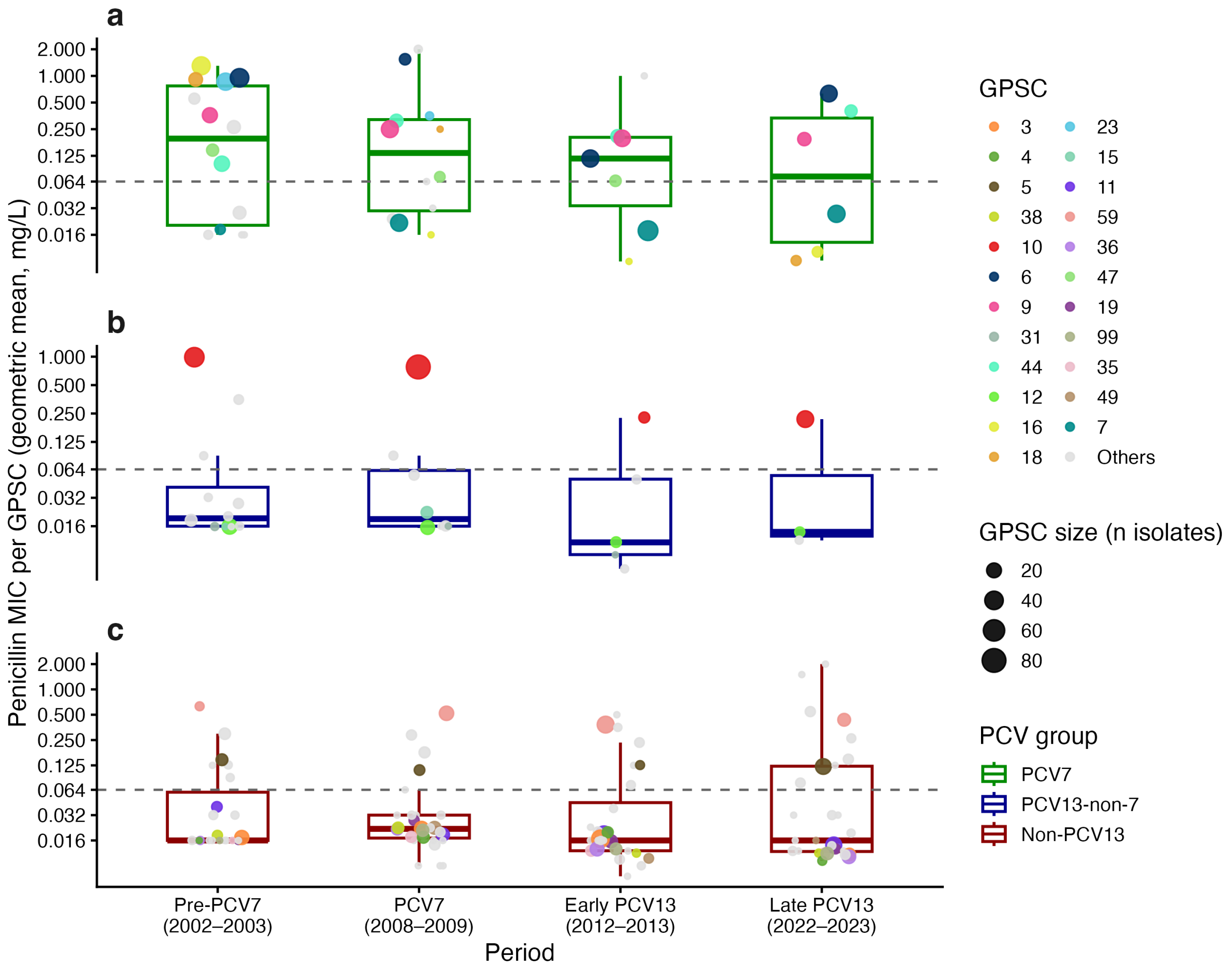


**Supplementary Fig. S8 | GPSC penicillin susceptibility across study periods**. **a–c**, For each period, points represent individual GPSCs plotted at the geometric mean MIC across isolates within that GPSC (mg/L; log scale). Point size is proportional to the number of isolates contributing to the GPSC estimate. Boxplots summarise the distribution of GPSC-level geometric mean MICs within each PCV group. The dashed horizontal line indicates the penicillin non-susceptibility threshold (MIC = 0.064 mg/L). Colours denote GPSC lineages (only high-frequency GPSCs are coloured and shown in the legend; low-frequency GPSCs are grouped as “Others” and coloured in grey).

### *Data-informed models for GPSC carriage rates over time*

**Supplementary Table S7 | Model coefficients for carriage rates of GPSC groups over study periods (2002–2023).**

| Model | Coefficient* | | | |
|---|---|---|---|---|
| Dataset | | **Estimate** | **Standard error** | ***P* value** |
| **Carriage rate£ of** | | | | |
| **PCV7-type GPSCs** | Intercept | -4.292 | 0.071 | <0.0001 |
| | Year | -0.263 | 0.046 | <0.0001 |
| | $Year^2$ | 0.214 | 0.051 | <0.0001 |
| | Mean log(MIC)# | -0.113 | 0.043 | 0.009 |
| **PCV13 non-PCV7-type GPSCs** | Intercept | -5.539 | 0.176 | <0.0001 |
| | Year | -3.835 | 0.487 | <0.0001 |
| | $Year^2$ | 0.028 | 0.078 | 0.727 |
| | $Year^3$ | 1.757 | 0.257 | <0.0001 |
| | Mean log(MIC)# | 0.365 | 0.073 | <0.0001 |
| **Non-PCV13-type GPSCs** | Intercept | -5.403 | 0.070 | <0.0001 |
| | Year | 0.324 | 0.056 | <0.0001 |
| | $Year^2$ | -0.176 | 0.052 | 0.0008 |
| | Mean log(MIC)# | -0.781 | 0.051 | <0.0001 |

£ Carriage rate is per 100 sampled children aged ≤2 years per year.
* Model coefficients are reported on the logit scale.
\# Mean log(MIC) corresponds to the mean log-transformed penicillin minimum inhibitory concentration of isolates within each GPSC in each period.
GPSCs were grouped according to their pre-PCV7 (2002–2003) serotype composition (>50%).
GPSC, Global Pneumococcal Sequence Cluster; IPD, invasive pneumococcal disease; PCV, pneumococcal conjugate vaccine; MIC, minimum inhibitory concentration.

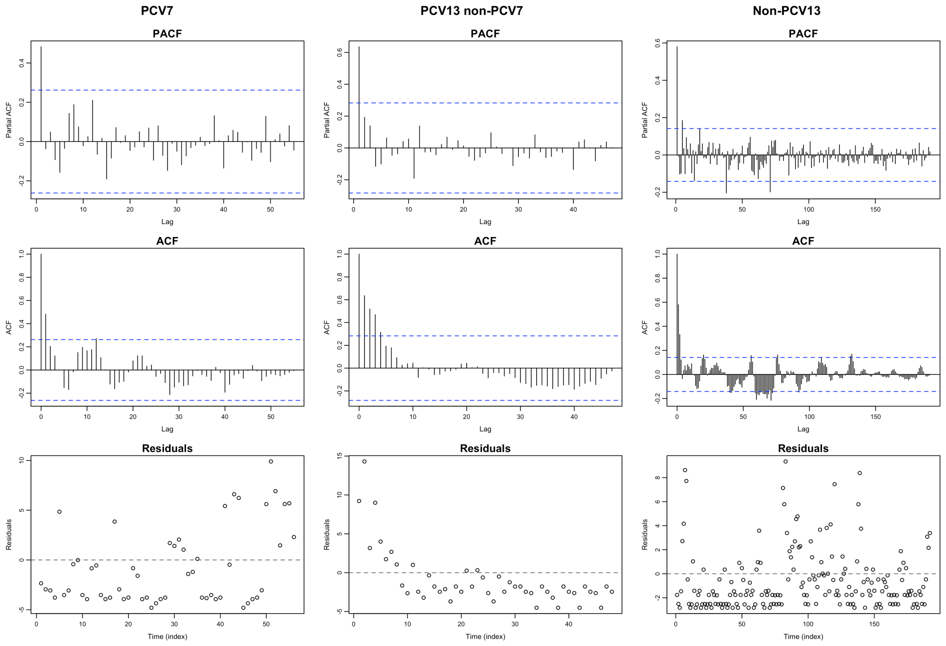


**Supplementary Fig. S9 | Goodness-of-fit diagnostics for the selected GPSC carriage data-informed models.** Residual diagnostics for polynomial logistic regression models for observed annual GPSC-group carriage rates in children aged ≤2 years. Columns correspond to PCV7, PCV13 non-PCV7 and non-PCV13 GPSC groups. Rows show partial autocorrelation functions (PACF), autocorrelation functions (ACF) and residual time series. Blue dashed lines indicate approximate 95% confidence bounds. Few temporal autocorrelation or systematic residual structure was observed.
PCV, pneumococcal conjugate vaccine**.**

### *Data-informed models for within-GPSC serotype composition over time*

**Supplementary Table S8 | Model coefficients for within-GPSC serotype composition of GPSCs in carriage over study periods (2002–2023).**

| Dataset | Coefficient* | Estimate | Standard error | *P* value |
|---|---|---|---|---|
| **Within-GPSC composition£ of** | | | | |
| **PCV7 serotypes** | | | | |
| | Intercept | -3.259 | 0.182 | <0.0001 |
| | Year | -1.620 | 0.110 | <0.0001 |
| | $Year^2$ | 1.062 | 0.125 | <0.0001 |
| **PCV13 non-PCV7 serotypes** | | | | |
| | Intercept | -2.064 | 0.164 | <0.0001 |
| | Year | -3.98 | 0.472 | <0.0001 |
| | $Year^2$ | -0.600 | 0.110 | <0.0001 |
| | $Year^3$ | 1.675 | 0.252 | <0.0001 |
| **Non-PCV13 serotypes** | | | | |
| | Intercept | 1.410 | 0.098 | <0.0001 |
| | Year | 1.815 | 0.101 | <0.0001 |
| | $Year^2$ | -0.528 | 0.094 | <0.0001 |

£ Within-GPSC serotype composition is the percentage of PCV7 (4, 6B, 9V, 14, 18C, 19F, and 23F), PCV13 non-PCV7 serotypes (1, 3, 5, 6A, 7F, and 19A), and non-PCV13 serotypes (serotypes not included in PCV13, including non-typeable isolates) within each GPSC per study period.
* Model coefficients are reported on the logit scale.
GPSC, Global Pneumococcal Sequence Cluster; PCV, pneumococcal conjugate vaccine.

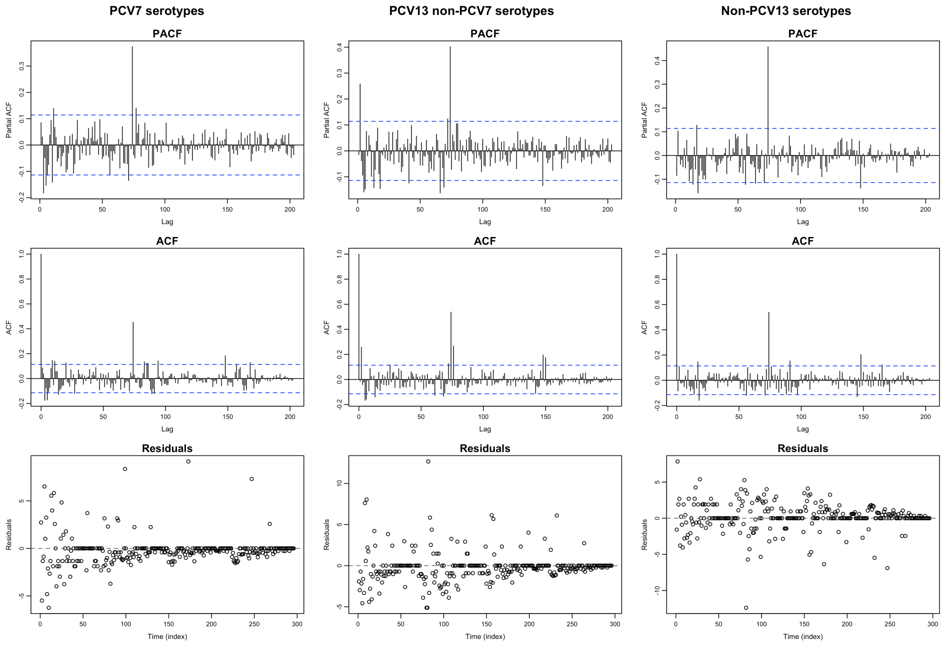


**Supplementary Fig. S10 | Goodness-of-fit diagnostics for the selected within-GPSC serotypes.** Residual diagnostics for polynomial logistic regression models for the proportion of serotype groups within each GPSC in carriage in children aged ≤2 years. Columns correspond to PCV7, PCV13 non-PCV7 and non-PCV13 serotype groups. Rows show partial autocorrelation functions (PACF), autocorrelation functions (ACF) and residual time series. Blue dashed lines indicate approximate 95% confidence bounds. No substantial temporal autocorrelation or systematic residual structure was observed.

### Supplementary Results 5 | Models for counterfactual GPSC carriage rates under serotype-fixed and lineage-fixed scenarios.

**Supplementary Table S9 | Model coefficients for carriage rates of counterfactual GPSC groups under the serotype-fixed and lineage-fixed scenarios across study periods (2002–2023).**

| Scenario/ dataset | Coefficient* | Estimate | Standard error | *P* value |
|---|---|---|---|---|
| **Serotype-fixed scenario** | | | | |
| **Carriage rate£ of** | | | | |
| **PCV7-type GPSCs** | | | | |
| | Intercept | -7.578 | 0.278 | <0.0001 |
| | Year | -1.863 | 0.165 | <0.0001 |
| | $Year^2$ | 1.134 | 0.188 | <0.0001 |
| | Mean log(MIC)$ | 0.679 | 0.082 | <0.0001 |
| **PCV13 non-PCV7-type GPSCs** | | | | |
| | Intercept | -5.38641 | 0.168 | <0.0001 |
| | Year | -4.046 | 0.462 | <0.0001 |
| | $Year^2$ | -0.673 | 0.165 | <0.0001 |
| | $Year^3$ | 1.412 | 0.263 | <0.0001 |
| | Mean log(MIC)$ | 0.346 | 0.072 | <0.0001 |
| **Non-PCV13-type GPSCs** | | | | |
| | Intercept | -3.788 | 0.046 | <0.0001 |
| | Year | 0.674 | 0.052 | <0.0001 |
| | $Year^2$ | -0.399 | 0.046 | <0.0001 |
| | Mean log(MIC)$ | -0.170 | 0.034 | <0.0001 |
| **Lineage-fixed scenario** | | | | |
| **Carriage rate£ of** | | | | |
| **PCV7-type GPSCs** | | | | |
| | Intercept | -3.574 | 0.035 | <0.0001 |
| | Mean log(MIC)$ | 0.032 | 0.035 | 0.37 |
| **PCV13 non-PCV7-type GPSCs** | | | | |
| | Intercept | -4.230 | 0.051 | <0.0001 |
| | Mean log(MIC)$ | -0.044 | 0.052 | 0.392 |
| **Non-PCV13-type GPSCs** | | | | |
| | Intercept | -6.047 | 0.066 | <0.0001 |
| | Mean log(MIC)$ | -0.491 | 0.063 | <0.0001 |

£ Carriage rate is per 100 sampled children aged ≤2 years per year.
* Model coefficients are reported on the logit scale.
$ Mean log(MIC) corresponds to the mean log-transformed MIC of isolates within each GPSC in each period.
GPSCs were grouped according to their pre-PCV7 (2002–2003) serotype composition (>50%).
GPSC, Global Pneumococcal Sequence Cluster; PCV, pneumococcal conjugate vaccine; IPD, invasive pneumococcal disease; CI, confidence interval; MIC, minimum inhibitory concentration.

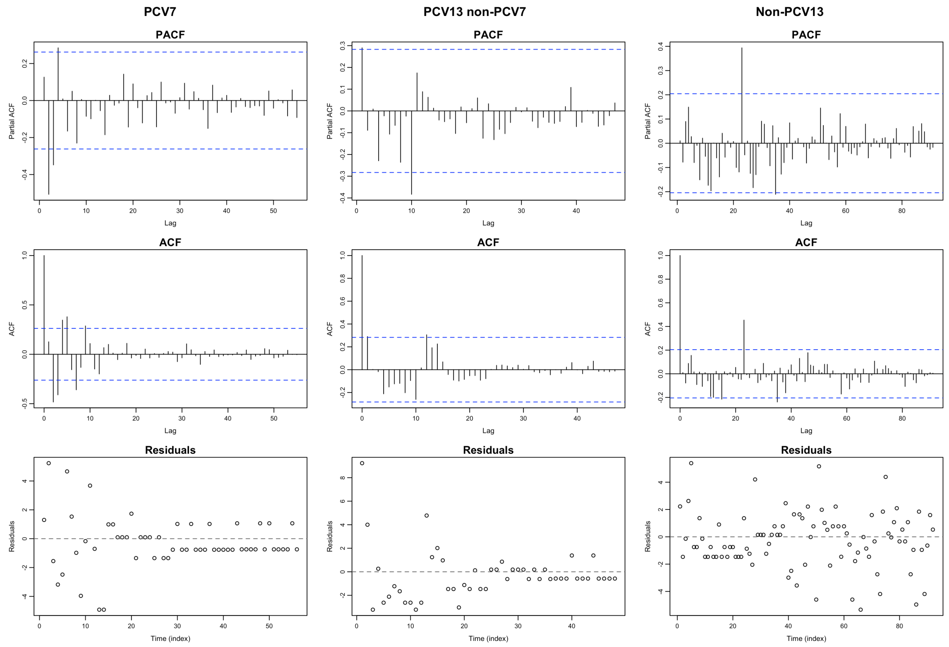


**Supplementary Fig. S11 | Goodness-of-fit diagnostics for the selected serotype-fixed scenario models.** Residual diagnostics for polynomial logistic regression models for the counterfactual annual carriage rate of GPSC groups in children aged ≤2 years under the serotype-fixed scenario. Columns correspond to PCV7, PCV13 non-PCV7 and non-PCV13 GPSC groups. Rows show partial autocorrelation functions (PACF), autocorrelation functions (ACF) and residual time series. Blue dashed lines indicate approximate 95% confidence bounds. No substantial temporal autocorrelation or systematic residual structure was observed.

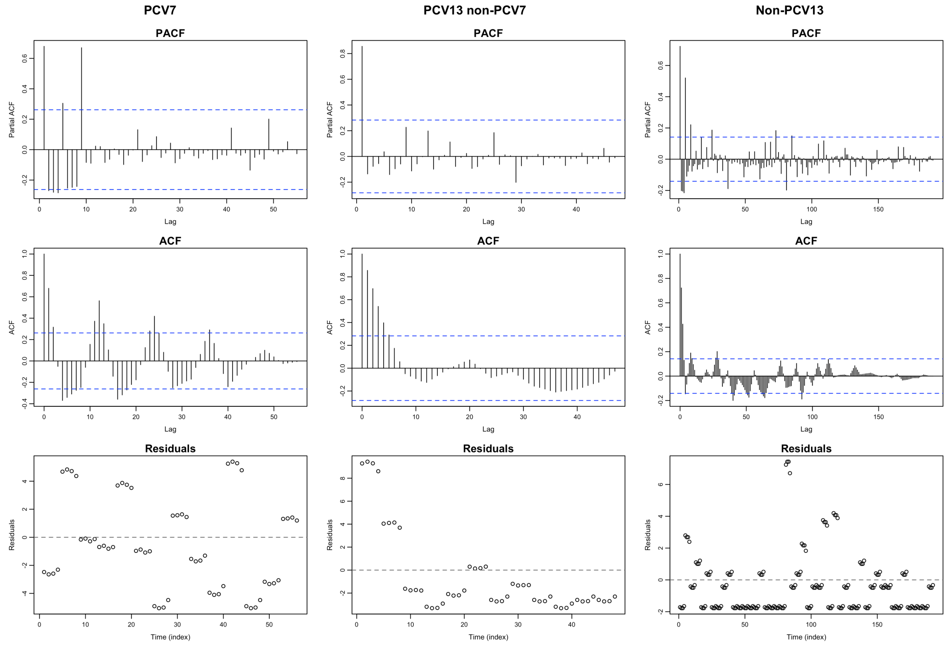


**Supplementary Fig. S12 | Goodness-of-fit diagnostics for the selected lineage-fixed scenario models.** Residual diagnostics for polynomial logistic regression models for the counterfactual annual carriage rate of GPSC groups in children aged ≤2 years under the lineage-fixed scenario. Columns correspond to PCV7, PCV13 non-PCV7 and non-PCV13 GPSC groups. Rows show partial autocorrelation functions (PACF), autocorrelation functions (ACF) and residual time series. Blue dashed lines indicate approximate 95% confidence bounds. Substantial temporal autocorrelation and residual structure was related to variability in individual GPSC distribution.

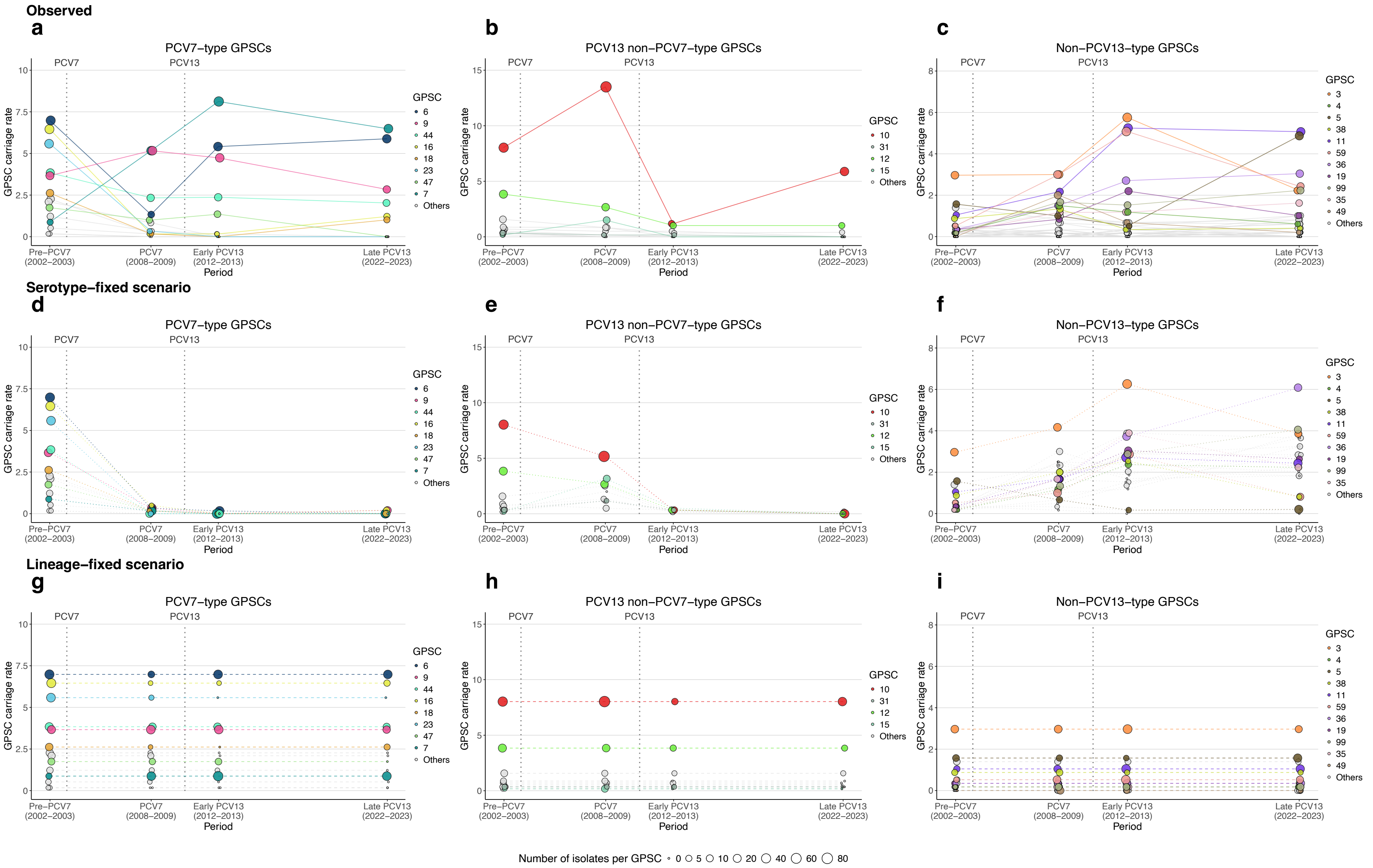
Observed
a
PCV7–type GPSCs
b
PCV13 non–PCV7–type GPSCs
c
Non–PCV13–type GPSCs
Serotype–fixed scenario
d
PCV7–type GPSCs
e
PCV13 non–PCV7–type GPSCs
f
Non–PCV13–type GPSCs
Lineage–fixed scenario
g
PCV7–type GPSCs
h
PCV13 non–PCV7–type GPSCs
i
Non–PCV13–type GPSCs
PCV7
PCV13
GPSC carriage rate
Pre–PCV7 (2002–2003)
PCV7 (2008–2009)
Early PCV13 (2012–2013)
Late PCV13 (2022–2023)
Period
GPSC
6
9
44
16
18
23
47
7
Others
GPSC
10
31
12
15
Others
GPSC
3
4
5
38
11
59
36
19
99
35
49
Others
Number of isolates per GPSC 0 5 10 20 40 60 80

**Supplementary Fig. S13 | Observed and counterfactual GPSC carriage rates. a–i**, Observed (**a–c**) and counterfactual GPSC annual carriage rates (per 100 sampled children aged ≤2 years) inferred under the serotype-fixed scenario (**d–f**), in which within-GPSC serotype composition was held constant over time, and under the lineage-fixed scenario (**g-i**), in which GPSC carriage rates were fixed across periods, across vaccine periods (2002–2023). GPSCs were grouped according to their pre-PCV7 (2002–2003) serotype composition (>50%) as PCV7-type (**a,d,g**), PCV13 non-PCV7-type (**b,e,h**), and non-PCV13-type (**c,f,i**). Each dot represents one GPSC, with dot size proportional to the observed number of isolates in each period. High-frequency GPSCs (>1% of isolates across the study period) are coloured and indicated in the legend; low-frequency GPSCs are shown in grey. Vertical dashed lines indicate the progressive implementation of PCV7 from 2003 and the introduction of PCV13 in 2010.
PCV, pneumococcal conjugate vaccine; GPSC, Global Pneumococcal Sequence Cluster.
